# Response of gut microbiota to feed-borne bacteria depends on fish growth rate: a snapshot survey of farmed juvenile *Takifugu obscurus*

**DOI:** 10.1101/2020.08.24.265785

**Authors:** Xingkun Jin, Ziwei Chen, Yan Shi, Jian-Fang Gui, Zhe Zhao

## Abstract

Understanding the ecological processes in controlling the assemblage of gut microbiota becomes an essential prerequisite for a more sustainable aquaculture. Here we used 16S rRNA amplicon sequencing to characterize the hindgut microbiota from cultured obscure puffer *Takifugu obscurus*. The gut microbiota is featured with lower alpha-diversity, greater beta-dispersion and higher average 16S rRNA copy numbers comparing to water and sediment, but far less so to feed. SourceTracker predicted a notable source signature from feed in gut microbiota. Furthermore, effect of varying degrees of feed-associated bacteria on compositional, functional and phylogenetic diversity of gut microbiota were revealed. Coincidently, considerable increase of species richness and feed source proportions both were observed in slow growth fugu, implying a reduced stability in gut microbiota upon bacterial disturbance from feed. Moreover, quantitative ecological analytic framework was applied and the ecological processes underlying such community shift were determined. In the context of lower degree of feed disturbance, homogeneous selection and dispersal limitation largely contribute to the community stability and partial variations among hosts. Whilst with the degree of feed disturbance increased, variable selection leads to an augmented interaction within gut microbiota, entailing community unstability and shift. Altogether, our findings illustrated a clear diversity-function relationships in fugu gut microbiota, and it has implicated in a strong correlation between feed-borne bacteria and host growth rate. These results provide a new insight into aquaculture of fugu and other economically important fishes, as well as a better understanding of host-microbe interactions in the vertebrate gastrointestinal tract.

**IMPORTANCE:** Environmental bacteria has a great impact on fish gut microbiota, yet little is known as to where fish acquire their gut symbionts, and how gut microbiota response to environmental bacteria. Through the integrative analysis by community profiling and source tracking, we show that feed-associated bacteria can impose a strong disturbance upon fugu gut microbiota. As a result, marked alterations in the composition and function of gut microbiota in slow growth fugu were observed, which is potentially correlated with the host physiological condition such as gastric evacuation rate. Our findings emphasized the intricate linkage between feed and gut microbiota, and highlighted the importance of resolving the feed source signal before the conclusion of comparative analysis of microbiota can be drawn. Our results provide a deeper insight into aquaculture of fugu and other economically important fishes, and have further implications for an improved understanding of host-microbe interactions in the vertebrate gastrointestinal tract.

## 1. INTRODUCTION

Microbiome can directly or indirectly affect the host’s physiological functions such as metabolism, immune defense, growth and development, as evidenced by increasingly large body of microbiome researches (1). Integrative conceptual frame of holobiont had been proposed, in which animals and plants are no longer seen as autonomous entities but rather as biological networks comprised with both host and its associated microbiome (2–4). Understanding the factors that underlie changes in such interactive network is essential, and could provide further insights into how the normal balance between host and microbiome (eubiosis) can be maintained or intervened if disrupted (dysbiosis)(1, 5). Systematically surveys of a broad range of environments and host species using the contemporary high-throughput sequencing technologies, have unravelled the structural and functional diversity of complex microbial communities and enabled indepth studies of microbiome-associated ecology, physiology and molecular function (6–8).

Exploring the composition and function of gut microbiome on fish have evoked increased research interest, not only for its representativeness as the most diverse vertebrate lineage, they also have a great economic significance, particularly in aquaculture (9). The development of high-potential microbiome-based innovations offers opportunities for disease control and health promotion in aquaculture practice (10, 11). Therefore, there is an urgent need of establishing an optimized microbial management strategy to achieve a more sustainable aquaculture (12). As a prerequisite, understanding the ecological processes and mechanisms in controlling the assemblage of fish associated microbiome becomes critical (13, 14).

Fish gut can be considered as a dynamic ecosystem in which spatial-temporal disturbance in the microbial nutrient niches shape the bacterial community (15, 16). The potential niche space in fish gut environment can be determined by the host in multiple ways (17), yet it also can be largely determined by the diet as well as it’s associated bacteria (18–20). Recent findings suggest that a limited numbers of bacterial taxa are shared inter-individually from a given host population and comprise a core gut microbiota (21–23). Fish-associated microbiota can be potentially acquired from a variety of sources with which organisms come into intimate biological interactions (17).

*Takifugu* (aka fugu), a genus of pufferfish (*Tetraodontidae*), is better known to be able to accumulate a potent neurotoxin, tetrodotoxin, and possess high toxicity (24). As a vertebrate, fugu displays several peculiarities in its genome, such as shortened intergenic and intronic sequences devoid of repetitive elements (25, 26), thereby makes it distinctive animal model for genome evolution study (27). Moreover, the artificially reared fugu becomes non-toxic and is considered a delicacy and commercially farmed widely across many East Asian countries including Japan, Korea and China (28, 29). In 2016, China had deregulated the fugu farming industry, and large-scale aquacultre of two fugu species *T. rubripes* and *T. obscurus* were licensed. With the market demand continues to grow, the aquaculture fugu production is progressively increased (24, 28). Therefore, understanding how and determining the extent to which the rearing environments effect fugu health and production is needed.

As a demersal fish, fugu live on or near the bottom of water, foraging food by scraping them off the surface (30). Meanwhile, feces from farmed fugu can also play a part of bacterial enrichment in rearing water and sediment as seen in other fish species (31, 32). Such dynamic reciprocal interactions might be expected to lead to a closer association between the gut of farmed fugu and its rearing environment to some extent, i.e. distinctions in the gut bacterial communties mirrored the environments and *vice versa*. Especially while considering the abundant bacterial species in the surrounding environments as a potential reservoir (19, 33, 34). On the other hand, unlike the pond water and sediment, the feed and gut are inherently linked by the high nutrient niche, which might impose a strong environmental selection of microbiota with functional traits linked to copiotroph (13, 35). Despite the evidences for the broad existence of core gut microbiota, accumulating studies had also shown substantial inter-individual variation in the gut microbial communites for both cultured (9, 36) and wild-captured fish species (37–39). Several factors can contribute to such variability, including environmental filtering either by the host, e.g. variations between regions of the gastrointestinal tract, state of health and nutrition; or by the rearing environments such as diet (40, 41). Such variations should be resolved before the conclusion of comparative analysis of microbiota can be drawn.

Environmental bacteria has a great impact on fish gut microbiota (12, 32, 33), yet little is known as to where fish acquire their gut symbionts, and to what extent gut microbiota response to it. Here we used 16S rRNA amplicon sequencing to characterize the hindgut microbiota from cultured obscure puffer (*T. obscurus*) in comparions with its rearing environment. Juvenile offsprings from a full-sib family were reared in a closed indoor aquaculture facility with consitent pond management and husbandry condition as well as feeding procedure. We shown that farmed fugu had a distinct gut microbiota with markedly lower alpha-diversity and higher beta-dispersion in comparison to its rearing environment including feed, water and sediment. Furthermore, effect of varying degrees of feed-associated bacteria on compositional, functional and phylogenetic diversity of gut microbiota were revealed. Furthermore, implications of fish growth-dependent community-level responses to feed-associated bacteria and the underlying relative importance of ecological processes were discussed. Our findings illustrated a clear diversity-function relationships in fugu gut microbiota, and emphasized the intricate linkage between feed and gut microbiota. These results provide a deeper insight into aquaculture of fugu and other economically important fishes, and have further implications for an improved understanding of host-microbe interactions in the vertebrate gastrointestinal tract.

## 2. RESULTS

In this study, the microbiota from fugu gut and its rearing environments were profiled using 16S rRNA gene amplicon sequencing. Using Illumina MiSeq, we obtained a total of 2.6 million reads with an average length of 410 bp from 82 samples (fish=61; feed=3; water=9; sediment=9). After quality filtering and chimeras removal using USEARCH followed by taxonomy-based filtering of plastids (chloroplast) sequences via RDP classifier, a total of 2,289 exact sequence variants (ESVs) were clustered. A total of 2.2 million high-quality reads were retained (mappable to ESVs) with average 26,712 reads per sample (ranging from 7,244 to 66,203; Table S1). Further, datasets were rarefied at 7,000 sequences per sample, followed by coverage estimates by rarefaction curves to ensure a sufficient sequencing depth and to include all samples for downstream analysis (Figure S1).

### 2.1. Comparative analysis of microbiota shows distinctions between fugu gut and rearing environment

The calculated indices of Richness (Observed ESVs), Chao1, Shannon and Simpson, congruously, indicated that the alpha-diversity of fugu gut microbiota was markedly lower comparing to that of rearing environment including feed, water and sediment (Figure S2). In addtion, statistical analysis of beta diversity based on Bray-Curtis distance shown that the pond (n=3) does have certain effect on community dissimilarity (R2=0.11; P=0.001; MeanSqs=0.97; Table S2), thereby ‘pond’ was set as a strata. Further, the community composition was revealed to be significantly differed among sample groups (PERMANOVA, R2=0.24; P=0.001; MeanSqs=2.08; strata=‘pond’; Figure. 1A). In addition, the permutational analysis of multivariate dispersion (PERMDISP) results showed that the sample groups differed in homogeneities, and the intra-group beta dispersion was significantly lower in feed group than any other groups (TukeyHSD, P<0.05, Table S2), possibly due to the lower samples size of feed (n=3) (Figure S3). Moreover, the abundance-weighted UniFrac distances of all intra-group pairs were significantly shorter than that of inter-group pairs (P<0.001;Mann–Whitney U), amongst others the fugu gut had the greatest within-group dispersions (F.F: range 0.069 ~0.996; coefficient of variation, 0.340) suggesting a much greater variability (Figure 1B).

**FIG 1.**
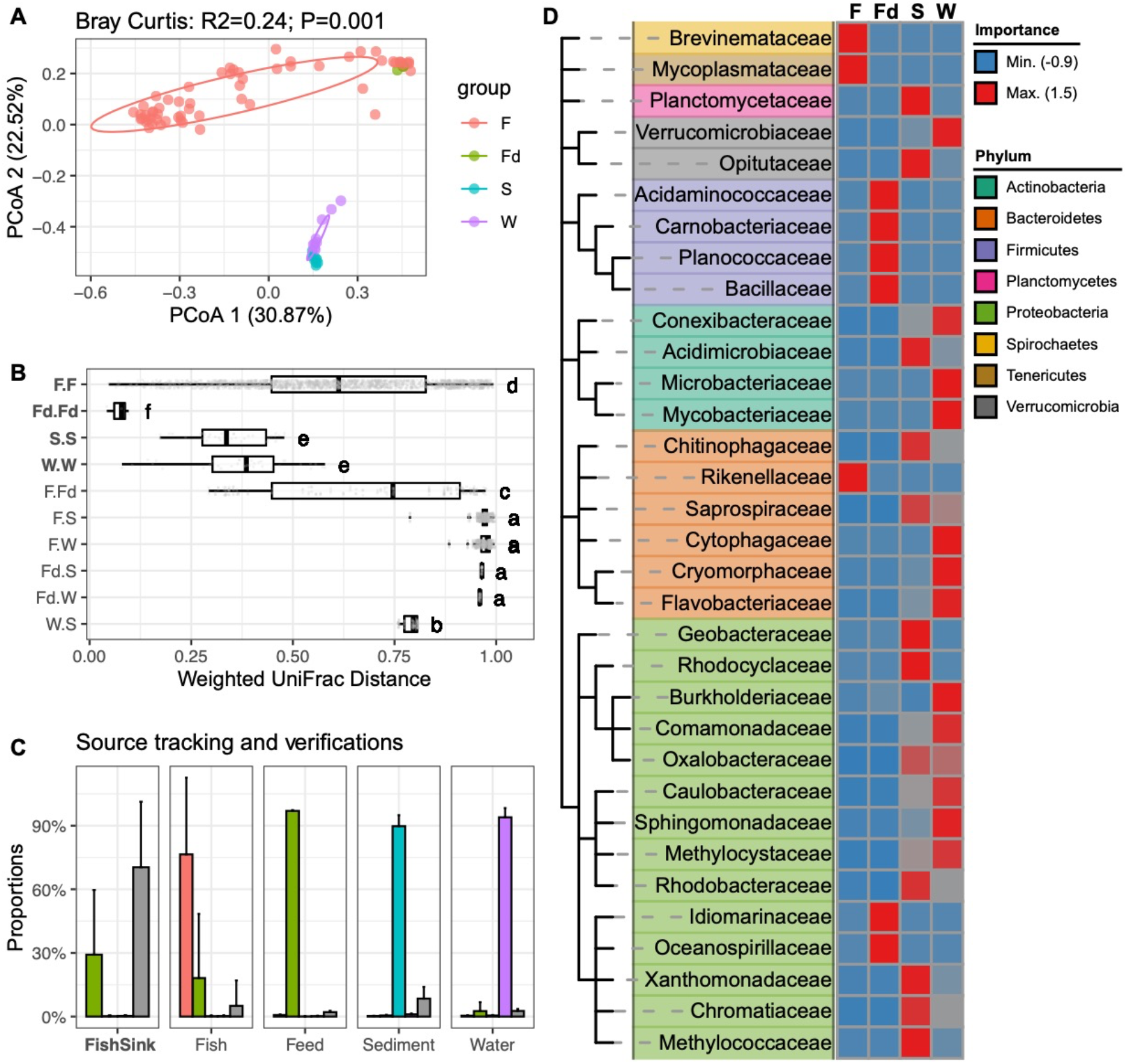
Comparions of different bacterial communities within the fugu rearing ecosystems. **(A)** Two-dimensional scatter plot of all bacterial communities based on Principal coordinate analysis (PCoA) on Bray-Curtis dissimilarity. Percentage of explained variance and statistical significance were reported by PERMANOVA. A normal data ellipse for each group was drawn at confidence level of 0.68. Sample groups: ‘F’: Fugu gut; ‘Fd’: feed; ‘S’: sediment; ‘W’: water. **(B)** Boxplot shows the intra-/inter-groups pairwise comparisons of weighted UniFrac distances across all sample groups. For each box, the vertical bold bar denote medians; the width of box denotes the interquartile range (25th percentile~ 75th percentile); the whiskers mark the values range within 1.5 times interquartile. Lower-cased letters denote statistical significance reported by Mann–Whitney U test at confidence level of 0.95. **(C)** Source tracking of gut microbiota from rearing environments. The far left facet shows the SourceTracker-estimated proportions of gut microbiota from its surrounding environment; the righter four facets respectively show the verifications using one each of the four sample groups as ‘source’. Bars denote average proportions of each source for a indicated community as sink (X-axis) using trained source samples (F=61, Fd=3, S=9, W=9) rarafied by 1000 ESVs. Error bars represent standard deviations. Bars were coloured according to the indicated sources, of which ‘Unknown’ represent source unidentified from trained source data. Color code is consistent with Figure 1A excepting the ‘unknown’ source was shown in grey. **(D)** Random-Forest identified bacterial families that accurately discriminated bacterial community groups. Featured taxa were hierarchically clusterd according to NCBI taxonamy, and were color-coded by phylum ranks. Heatmap shown the centered classific importance of each indicated taxa (row) in discriminating a given bacterial community (colomn) from others.

We further estimated the proportions of feed, sediment and water as a potential environmental bacteria ‘source’ for the gut microbiota (as ‘sink’) using SourceTracker. Results shown that the majority of bacterial proportions in fugu gut microbiota were, in fact, not identifiable from any given source (‘unknown’, 70.42 % ± 30.87% s.d.), suggesting a higher degree of host specificity. Yet, a notable proportion was predicted from feed (29.19% ± 30.53% s.d.)(far left facet “FishSink” in Figure 1C and Table S3). Moreover, verifications using one each of the four sample groups as ‘source’ had confirmed the accuracy of the predictive measures in classifying among different groups, despite a minor proportion from an unidentified source (unkown) within each group (four right-side facets in Figure 1C).

The characteristic taxa were classified as a attribute in distinguishing among different groups of bacterial community using Random-Forest machine-learning approach. Results post ten-fold cross-validation shown that at least 34 bacterial families can be used to classify all four sample groups with 1.22% error rate (Figure S4, Table S4). Of these, 33 families were taxonomically categorized into eight phylums according to the NCBI Taxonamy (Figure 1D). Three indicative bacterial families for fugu gut microbiota were identified, namely *Brevinemataceae (Spirochaetes), Mycoplasmataceae (Tenericutes)* and *Rikenellaceae* (*Bacteroidetes*). Whereas, greater numbers of diverse indicative families for environmental sample groups were determined, and brought into correspondence with the observed high bacterial diversity in environmental microbiota.

### 2.2. Community structure in fugu gut are correlated with feed source signial and host body mass

Individual fugu were clusted based on unsupervised k-means computed from body mass measures including weight (unit: gram) of body, liver and gonad, as well as fork length (unit: centimeter). Using gap statistic method, the optimal clustering numbers was determiend to be 3, which we defined as ‘fast’ (n=24), ‘medium’ (n=20) and ‘slow’ (n=17) (Figure 2A and Figure S5). The quality of representation (cos^2^) for body weight, liver weight, fork length and gonad weight on the first principal component (PC1, 88.1% of total variations) were 0.96, 0.93, 0.92 and 0.72, respectively (Table S2). The PC1 value was used as a proxy for individual fish body mass for downstream analysis.

**FIG 2.**
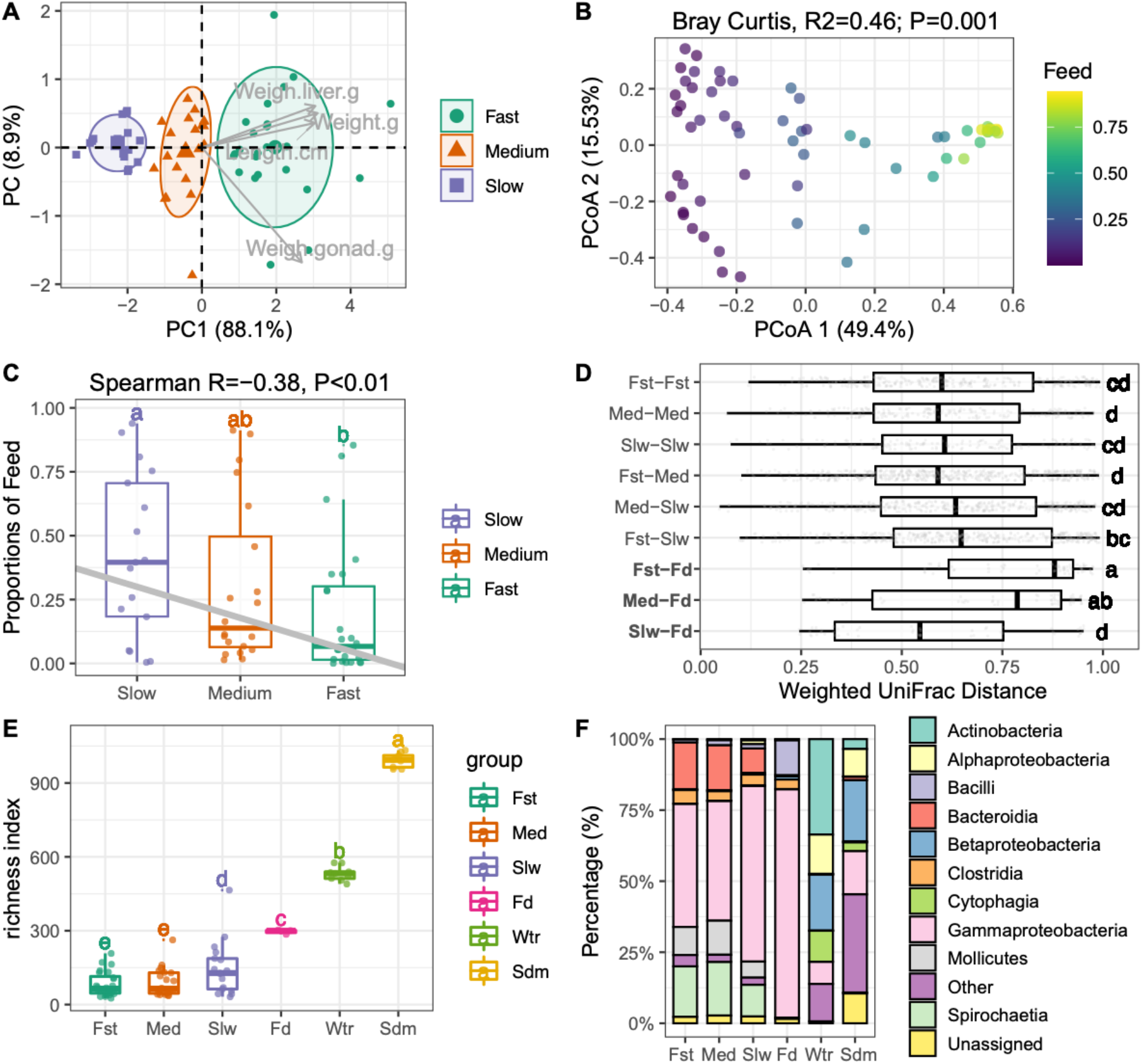
Community structure of fugu gut microbiota are correlated with feed source signial and host body mass. **(A)** Two-dimensional scatter plot of all fish individual based on Principal Components Analysis (PCA) of four body mass metrics. The grey arrow points shown the correlated body mass measures. Three optimal groups namely “fast”, “medium” and “slow” were identified by K-mean clustering followed by Gap statistic. A normal data ellipse for each group was drawn at confidence level of 0.68. **(B)** Principal coordinate analysis of gut microbiota based on Bray-Curtis distance. Percentage of explained variance and statistics (PERMANOVA test with 999 permutations) were shown as figure title. Each community point was coloured by the proportion of feed source signal. **(C)** SourceTracker-estimated proportions of feed bacteria for each gut community grouped by host body mass. Grey line denotes the linear regression of feed proportions and fugu body mass (Spearman’s Rank), stataistics was indicated in figure title. **(D)** Boxplot of group-paired weighted UniFrac distances showing the dissimilarities between gut and feed were reduced in slow growth fugu (Slw-Fd) comparing to fast growth one (Fst-Fd). **(E)** Species richness (Numbers of oberseved ESVs) among six groups of bacterial community. For each box in (**C~E**), the bold bar denote medians; the height of box denotes the interquartile range (25th percentile~ 75th percentile); the whiskers mark the values range within 1.5 times interquartile. Lower-cased letters denote statistical significance reported by Mann–Whitney U Test at confidence level of 0.95. **(F)** Stackbars showing the relative abundance of top-10 bacterial classes across six groups of community. Bacterial class which has a lower relative abundance were grouped into “Other”. The detailed results of replicated samples were shown in Figure S7.

Statistical analyses of beta diversity were performed to infer the potential factors underlying the large inter-individual beta-dispersion of gut microbiota. PERMANOVA analysis shown feed source signals was the best predictor of community composition (R2=0.46; *P*=0.001; MeanSqs= 6.68; strata:Pond; Figure 2B), followed by fish body mass (R2=0.07; *P*<0.01; MeanSqs=0.98; strata:’Pond’). Importantly, negative correlation between the fish body mass and the feed source signal was observed (Spearman’s Rank R=-0.38; *P*<0.01, Figure 2C). Moreover, the feed source proportions were significantly enriched in slow growth fish (42.2%) comparing to fast one (19.3%) (TukeyHSD, *P*<0.05). These results suggest the varing degree of effect of feed-associated bacteria on gut microbiota is dependent on host growth rate. Specifically, the gut community dissimilarity distances to feed for slow growth fish were significantly shorter than that for fast one, as evidenced by the comparisons of abundance-weighted UniFrac distances (*P*<0.001;Mann–Whitney U, Figure 2D).

The estimated indices of alpha-diversity were positively correlated with feed source signals, but negatively correlated with fugu body mass (Figure S6). Futher, markedly increased richness of gut microbiota from slow growth fish was observed comparing to fast group (*P*<0.05; Mann–Whitney U, Figure 2E). The taxonomic compositions of bacterial communities were differed among sample groups (Figure S7; Table S5). In the bacterial class level (Figure 2F), *Actinobacteria, Alpha-, Beta-, Gamma-proteobacteria* and *Cytophagia* were commonly found to be dominant in both water (sum 85.9%) and sediment (sum 53.0%), whereas *Actinobacteria* and *Cytophagia* were more present in water (33.6% and 10.9%) than to sediment (3.5% and 3.0%). Feed associated bacterial communities were mainly composed of *Gamma-proteobacteria* (80.4%), *Bacilli* (12.2%) and *Clostridia* (3.4%). The gut bacterial communities of the three fish clusters were mainly composed of *Gamma-proteobacteria* (avg. 49.1%), *Spirochaetia* (avg. 16.0%), *Bacteroidia* (avg. 13.7%), *Mollicutes* (avg. 9.2%) and *Clostridia* (avg. 4.0%). Moreover, the fish gut-specific bacterial classes including *Bacteroidia, Mollicutes* and *Spirochaetia* were more present in fast and medium groups, whereas *Gamma-proteobacteria* was more present in the slow group.

LEfSe algorithm was used to identify differentially abundant bacterial taxa that are corresponding with the differences between fugu growth rate. A total of 53 discriminant bacterial clades seperating three fish clusters with statistical significance (Kruskal-Wallis sum-rank, *P*<0.05) were detected (Figure S8 and Table S6). The overall characteristic bacterial families were more detected in slow group, which is in consistent with its observed high species richness. Amongst others, *Pseudomonadaceae* was the most pronounced bacterial family (log_10_-LDA Scores >5) that were overrepresented in slow group.

### 2.3. Functional shifts of fugu gut microbiota in response to feed-borne bacteria

Understanding the phenotypes and functional capacities of the microbes within a community is critical to determine the composition-function relationships. To this end, we used PICRUSt and BugBase respectively to infer the functional profiles and organism-level phenotypes of different microbial communities based on 16S rRNA gene. Increased ribosomal content boosts more protein translation and stronger metabolism, thereby the number of rRNA genomic copies is positively correlated with bacterial growth rate (42), and can be used as a proxy for phenotypically distinguishing between copiotroph and oligotroph (43, 44). Using the precalculated 16S rRNA counts by PICRUSt, the avarage 16S rRNA copy number per sample group were estimated. The results shown that bacterial communities from fugu gut (3.6) and feed (3.4) had greater average copy numbers than that from water (2.0) and sediment (1.9), indicating the copiotrophic dominance communities in both gut and feed (Figure 3A).

**FIG 3.**
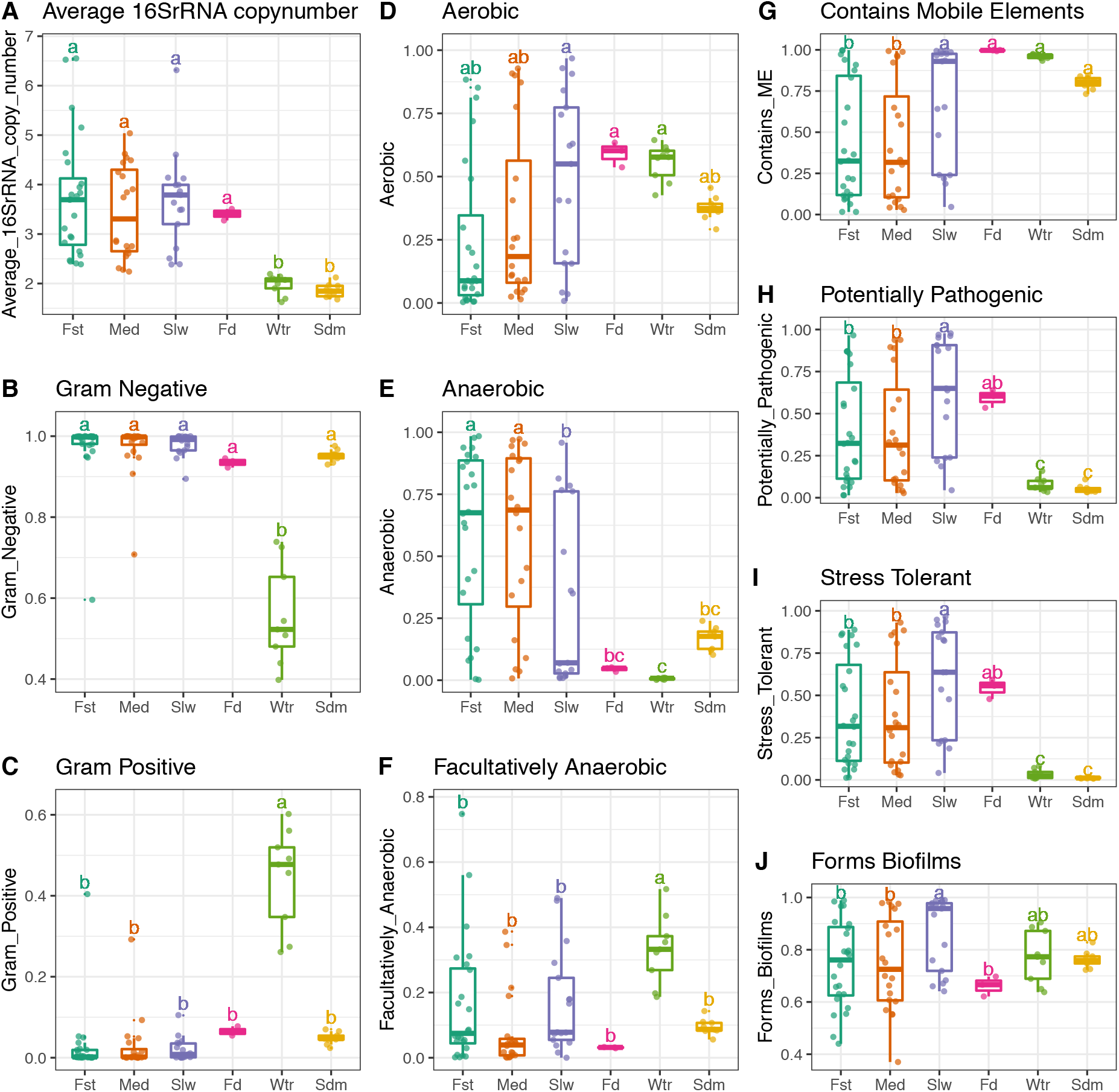
Phenotype inference of bacterial communities from the fugu rearing ecosystems. Estimatied relative abundances for each indicated bacterial phenotype was compared across different sample groups. **(A)** Averaged 16S rRNA copy numbers were compared among sample sites of bacterial communities. Points denote the mean copy number calculated from a given bacterial community. The following organism-level phenotypes were infered by BugBase. Relative abundances of bacteria differing in Gram staining were shown in **(B)** for Gram Negative and **(C)** for Gram Positive. Relative abundances of bacteria differing in oxygen tolerance phenotypes were shown in **(D)** for Aerobic, **(E)** for Anaerobic and **(F)** for Facultatively Anaerobic. Relative abundances of bacteria differing in latent pathogenicity phenotypes were shown in **(G)** for containing mobile elements, **(H)** for potentially pathogenic, **(I)** for oxidative stress tolerance and **(J)** for biofilm formation. See Figure S9 for the related taxa contributions of the relative abundance of bacteria possessing each phenotype. For each box, the horizontal bold bar denote medians; the height of box denotes the interquartile range (25th percentile~ 75th percentile); the whiskers mark the values range within 1.5 times interquartile. Lower-cased letters denote groups and statistical significance reported by pair-wise Mann-Whitney U tests with false discovery rate correction at confidence level of 0.95.

Further, the phenotypical charasteristics including oxygen tolerance, Gram staining and pathogenic potential of bacterial communities were predicted by BugBase algorithm. It infered the gut most significantly to be the negative bacteria dominance communities (97%), followed by feed (93%) and sediment (95%) (Figure 3B). Whereas water microbiota was predicted to be comprised of both Gram negative (56%) and Gram positive (44%) bacteria (Figure 3B and C, and Figure S9). Regarding oxygen tolerance traits (Figure 3 D~F), gut was shown to be significantly dominant by anaerobic bacteria (51%) comparing to all the other environments (<20%), where is mostly dominant by aerobic bacteria as for feed (59%), or by both aerobic and facultatively anaerobic bacteria as for water (56% and 33%). Whereas such dominance was significnatly decreased as the proportions of aerobic bacteria increased in slow growth fugu (49%) comparing to both fast (24%) and medium (34%) groups. In deed, the infered proportions of anaerobic bacteria in gut microbiota is negatively correlated with feed source signals (Spearman’s Rank R=-0.79; *P*<0.01), but positively correlated with body mass (R=0.32; *P*<0.05, Figure S10). The opposite trends were observed for infered aerobic bacterial proportions. These results suggest the feed source signals is the most pronounced factor in contributing such phenotypic changes in gut microbiota, and it is depended on host body mass.

BugBase aslo predicted the fugu gut microbiota to have significantly fewer bacteria that contain mobile elements and more bacteria that are potentially pathogenic and oxidative stress tolerant than water and sediment (*P*<0.05, Mann–Whitney U, Figure 3 G~I). Further, bacteria that can form biofilms was not significantly different comparing fugu gut to both water and sediment (Figure 3J). Within gut microbiota, the bacterial proportions related to these four phenotypes were all positively correlated with feed source signals (R>0.7; *P*<0.05) but negatively correlated with body mass (R<-0.3, *P*<0.05, Figure S10). Altogether, these results have further emphasized that the functional changes in fugu gut microbiota is drived largely by feed-associated bacteria, and depended on host growth rate.

### 2.4. Ecological processes underlying bacterial community shift in fugu gut microbiota

To infer the underlying ecological mechanisms governing the process of bacterial community assemblage in fugu gut, the phylogenetic signals of gut bacterial communities were estimated using Mantel’s correlogram (45) and Blomberg’s K statistic (46), respectively. We found significant phylogenetic signal (*P*<0.05) correlating to changes of feed source signals and fish body mass, but only across very short phylogenetic distances (Figure S11). On this basis, we proceeded to quantify phylogenetic turnover among close-related bacteria (47). Considering the ecological influence on bacterial sub-community may varied with its relative abundance (48–50), the fugu gut microbiota (‘all’, 943 ESVs) were further partitioned into ‘abundant’ (abundance >0.1%, 63 ESVs) and ‘rare’ sub-community (abundance <0.1%, 880 ESVs), respestively. We concentrated on ‘all’, ‘abundant’ and ‘rare’ ESVs to extract phylogenetic distance from a given bacterial community. The results shown that the unweight standardized effect size of mean nearest taxon distances (ses.MNTD) for ‘all’, ‘abundant’ and ‘rare’ sub-communities from gut microbiota were consistently below zero (*P*<0.01, Figure S12), suggesting the importance of environmental filtering in driving gut bacterial communities to be more phylogenetically clustered. In addtion, the average ses.MNTD measures for abundant taxa were markedly higher than that of rare taxa (Figure S12), suggesting its lower degreed of phylogenetic relatedness.

Further we estimated the phylogenetic beta diversity (βMNTD) and in turn compared it against null model to estimated beta nearest taxon index βNTI (47, 51). The |βNTI| values > 2 indicate significantly phylogenetic turnover than expected. The Mantel’s test showed significant positive correlations between the inter-community βNTI values and the changes of feed source signals, for ‘all’, ‘abundant’ and ‘rare’ sub-communities, respectively (Figure 4A~C). These results suggest that the shift from homogeneous selection to variable selection in contributing to the gut bacterial community assemblage as changes of feed source signals increase.

**FIG 4.**
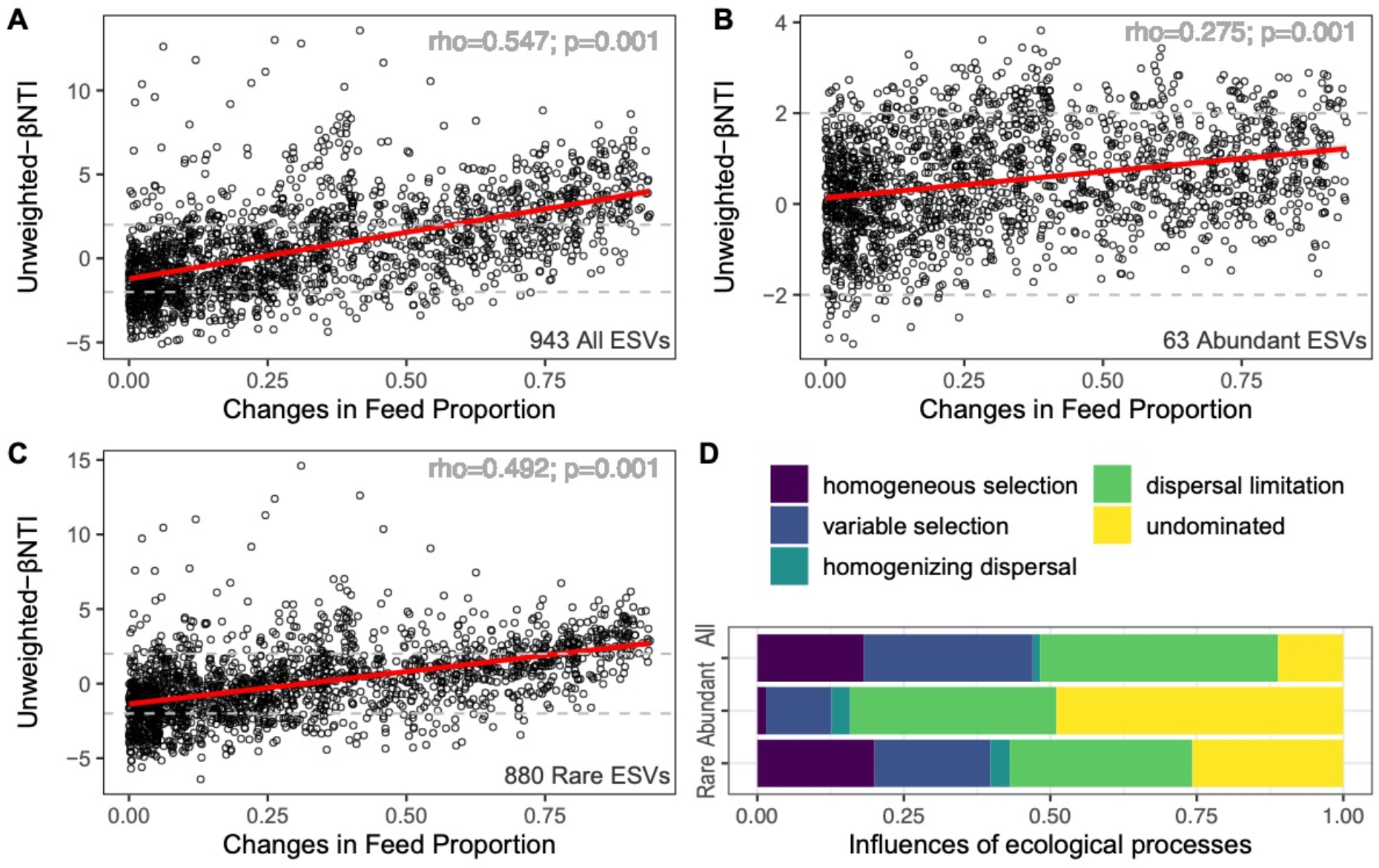
Influence of ecological processes in governing bacterial community assemblage in fugu gut. The correlations between phylogenetic beta diversity βNTI and differences in feed source signals in fugu gut microbiota for **(A)** all ESVs, **(B)** abundant ESVs sub-community and **(C)** rare ESVs sub-community. Red lines denote fitting of linear regressions and corresponding Spearman’s rank correlation coefficients were indicated in the upright corner within each subpanel. Grey horizontal dashed lines denote the cutoff of βNTI significance between −2 and +2. **(D)** Depicts the relative importance of different ecological processes in governing community turnover for ‘all’, ‘abundant’ and ‘rare’ sub-communities, respectively.

To quantify the ecological processes in communities that were not governed by selection (i.e. |βNTI| < 2), the number of these pairwise comparisons were fractionated considering |βNTI|<2 and |RC_bc_| < 0.95. The resulting fractions shown that dispersal limitations (40.6%) contribute largely in the process of community assemblage in fugu gut while considering all the taxa, followed by variable selection (28.9%) and homogeneous selection (18.1%). While comparing the sub-communities varied in taxa abundance, the relative importance of deterministic process (homogeneous and variable selection) was larger for ‘rare’ sub-community, whereas stochastic processes (dispersal limitation and ‘undominated’) was dominant in ‘abundanct’ sub-community (Figure 4D). In addtion, very few portions of homogenizing dispersal process were observed for all sub-communities. Furthermore, the neutral model explained a larger fraction of variation in the rare sub-community (R2=0.639,m=0.057) than the abundant one (R2=0.154, m=0.012), as well as increased migrations rate (Figure S13).

## 3. Discussion

Here we used 16S rRNA gene amplicon sequencing to explore the relations of bacterial communities between gut of cultured juvenile obscure puffer and its rearing environments. Our results shown that farmed fugu had a distinct gut microbiota with markedly lower richness (alpha-diversity) and higher dissimilarity (beta-dispersion) in comparison to its rearing environment including feed, water and sediment (Figure 1). Furthermore, effect of varying degrees of feed-associated bacteria on both composition and function of gut microbiota were revealed, and implications of fish growth-dependent community-level responses and underlying relative importance of ecological processes were found. These results (discussed below) provide a new insight into aquaculture of fugu and other farmed fishes that are of great economic importance, as well as a better understanding of host-microbe interactions in the vertebrate gastrointestinal tract.

### 3.1. Distinctions of fugu gut microbiota from its rearing environment

Herein, the fugu gut microbiota is characterized by lower species richness, higher beta-disperpsion and rRNA copy numbers in comparison to water and sediment, but far less so to feed (Figure 1 and 3), indicating a strong niche effect on gut microbiota along with considerable diet influence. Indeed, SourceTracker had predicted a notable source signature from feed (~30%) in contributing to fugu gut microbiota (Figure 1C), although the samples were obtained one day post-feeding. This could explain why the community composition differed largely between “gut & feed” and “water & sediment”, as the metabolic mode of copiotroph and oligotroph differed significantly (Figure 3A). Moreover, a larger proportion of ‘unknown’ source signature was also predicted, suggesting an uncharacterized bacterial source possibly (i) derived elsewhere or (ii) acquired before the sampling time. For the first scenario, it is less likely that farmed fugu had much contact with any other environmental source that had not been sampled, as the indoor pond managements and husbandry conditions both were identical within a single closed aquaculture facility. For the second scenario, several studies had pointed that the processes of bacterial colonization in fish gut, particularly during early development, are complex ever since the onset of spawning, and largely determined by the egg adhering microbiota and temporal rearing environment (17, 20, 34, 52). Therefore, it can be hypothesized that the early-life exposure to different bacterial communities as seen in other fishes, might also contribute to the assembly of fugu gut microbiota mentioned here.

Ringø and Birkbeck (53) proposed five essential criteria to be considered core (autochthonous) gut microbiota in fishes if they are (i) presented in healthy individuals; (ii) persistent during the whole lifespan of host; (iii) mutually presented in both wild and cultured populations; (iv) anaerobic; and (v) widely distributed in gastrointestinal tract. Our Random-Forest analysis identified three bacterial families, namely *Brevinemataceae, Rikenellaceae* and *Mycoplasmataceae*, as the most discriminative taxa for fugu gut (Figure 1D). In addition, Bugbase infered that both *Brevinemataceae* and *Rikenellaceae* contributed to the most proportions of anaerobic bacteria in fugu gut microbiota (Figure S9). Although literature concerning the microbiota of wild *T. obscurus* is scarce, recent studies shown that these bacterial families, particularly *Brevinemataceae* with greater dominance, can be also observed in many other wild *Takifugu* species, including *T. ocellatus*, *T. bimaculatus*, *T. xanthopterus* (54) and *T. niphobles* (55) regardless of methodological biases. It is remarkable since unlike these wild species that are native to salt and brackish waters, farmed *T. obscurus* was mainly reared in fresh water after hatching, with the only exceptions of spawning parent fishes and eggs that were transiently acclimated in saline water. Once again it demonstrates the potential influence of early-life environments in shaping the fugu gut microbiota. Altogether, these bacterial taxa meet most of the criteria to be considered core gut microbiota in fugu, yet further longitudinal experiments concerning multiple host developmental stages are warranted to completely define it.

### 3.2. Effects of feed-borne bacteria on gut microbiota differed among fugu at inconsistent growth rate

We next asked which factors (either intrinsic or extrinsic) potentially correlate with the large inter-individual beta-dispersion of gut bacterial community. Here, the gut contents were individually sampled from the hindgut of farmed fugu that were feed with identical commercial fish feed. In addition, the fish used here were offsprings from a full-sib family, thereby the potential genetic variations were kept to a minimum. Moreover, the environmental bacterial communities for both water and sediment were found to be consistent across different ponds (data not shown), thereby the likelyhood of such extrinsic variations could also be ruled out. In fact, we observed a large dispersion of body mass among the sampled fish population (Figure S5), by which they can be rigorously subdivided into three clusters, i.e. fast, medium and slow (Figure 2A). We shown that the feed source signals explained the most variations in the compositional difference in fugu gut microbiota, and its was negatively correlated with fish body mass (Figure 2B and C). Across the fishes clustered by body mass, both the composition and function of gut bacterial communities were found to be varied significantly, as evidenced by the comparative analysis of richness (alpha-diversity), dissimilarity (beta-diversity), discriminated abundant bacterial taxa (LEfSe) and orgnism-level phenotypes (BugBase) (Figure 2 and 3), respectively. These results illustrated a clear diversity-function relationships in gut bacterial communities, and more importantly, it implicated a strong correlation between gut bacterial communities and host growth rate. Specifically, markedly increased species richness driven by feed source proportions were observed in slow growth fugu, where a greater level of correlation between feed-associated bacteria and gut microbiota was implied.

Such coincidence of altered microbiota composition and function in the gut of slow growth fish were largely driven by a *gamma-proteobacterial* genus *Pseudomonas* (LEfSe, log_10_-LDA Score>5, *P*<0.05), notably a group of widespread aerobic bacteria found to be predominant in fish feed herein (>50%, Figure S7 and S9). Previous work had shown that *Pseudomonas* is highly persistent during long term storage of fish feed (56). Yet another study found its presence in water, contributed mostly by the fish feces, was positively associated with the dose of administrated commercial fish feed in a recirculating aquaculture system (31). Amongst others, *Pseudomonas* is the most frequently reported gut-associated bacteria across many different freshwater fish species (39). On the other hand, recent studies had also raised concerns that it might represent a common contaminating bacterial taxa in molecular biology reagents (57). Whilst, considering the very rare presence of *Pseudomonas* in all the water and sediment samples (Figure S7E), along with the negative amplification signal in our blank (no template) control experiment, it is highly unlikely that this bacteria taxa was introduced during libraries preparation as a contamination, rather than a biological clue reflecting the heterogeneous sample sources. Nevertheless, we could not explicitly estimate the extent to which the alteration of gut mirobiota was solely due to the introduced feed-borne bacteria, since the DNA-based amplicon sequencing, as was used here, is unable to ascertain (transcriptionally) active bacteria. Of note, dosens of ESVs filtered out as chloroplastid were mostly derived from the feed (~13.7% of total reads), but barely detected in gut (~0.1%). Such disproportion indicates that the overrepresent feed-borne bacteria in the gut of slow growth fugu were not exclusively due to the influx of DNA from dead or dormant cells (19). Rather, bacteria such as *Pseudomonas*, optimized to grow at high nutrient as copiotroph (with 6~10 average 16S rRNA copy number) (58), are known to be more competitive than oligotrophy in rapid growing or efficiently utilizing resource in the intestinal environment. Importantly, we found the response of individual gut microbiota to feed-borne bacteria varied with the fish growth rate, and became more pronounced in slow growth fugu.

Previous studies suggested that the transit rate and residence time of diet in the gastrointestinal tract may influence the gut microbiota and host-microbial interactions (16). Furthermore, the fish size was found to be negatively correlated with gastric emptying time, i.e. larger fish empty their stomachs at a faster rate than smaller ones within a consistent environment (59). And the fish gastric evacuation can be further enhanced by transgenetically overexpressions of host growth hormone gene (60). These findings offer support for our observations that proportions of feed source signals in gut microbiota was negatively correlated with fugu growth rate, which could be largely due to the inconsistent gastric evacuation rate among different sized hosts. Several studies had suggested that the feed associated microbiota could play a crucial role in host health and nutrition (18, 19, 61), yet little is know about the functional impact and the extent to which it exerted. Herein, BugBase predicted the gut of slow growth fugu (with higher feed source signal) to have significantly more bacteria that are potentially pathogenic and biofilms forming than other fish clusters (Figure 3). Such phenotypical differences are driven mostly by the higher proportion of both *Pseudomonadaceae* and *Enterobacteriaceae* (Figure S9), of which many members are generally considered commensals (61). But like other opportunistic pathogens, certain members from both bacterial families often contain genes for virulence factors and antibiotic resistance (62, 63). It’s interesting that, the *Pseudomonadaceae* that was considered as presumptive feed-borne bacteria had very high relative abundance in feed (65.9%); whereas *Enterobacteriaceae* is relatively rare in feed (0.3%) in comparison to gut (3.8% in slow growth fugu) (Figure S7). Therefore, we hypothesized that the observed alteration of fugu gut microbiota was determined by a combinational effects of (i) intestinal residence time allowing feed-borne bacteria to proliferate and (ii) response outcomes from interactions between allochthonous and autochthonous bacteria. The intuitive expection is that, in slow growth fugu, further reduced gastric evacuation rate and more inadequate anaerobic niche in intestinal lumen allowed the faster grow of aerobiotic feed-borne bacteria (mainly *Pseudomonas*), and simultaneously influenced the abundance of autochthonous bacteria by either enhancing or diminishing their growth. As an further extreme assumption, the increased feed-borne bacteria in gut microbiota may lead to an accumulation of negative effects (e.g. pathogenesis) for the host, thereby stimulated intestinal stress response and hampered metabolic and digestion in host can be expected, which consequecely could lead to a further reduce scope for the fish growth (lower fitness). Future studies concerning the ingested feed-borne bacteria and its interactions with both gut bacterial symbiosis and its host are critical for a better understanding of fish health and metabolism.

### 3.3. Quantitative estimation of ecological processes underlying the gut microbiota response to feed-borne bacteria

We found significant phylogenetic signals related to traits of feed source and body mass (*P*<0.05, Figure S11). In addtion, the mean within-community phylogenetic distances (ses.MNTD) in gut microbiota were all found to be negative (t-test, *P*<0.001), further comfirming all the ESVs were phylogenetically correlated more closely than expected by chance (Figure S12). Whereas, all the K values were estimated to be positive but closer to zero (Table S2), indicating a random or convergent pattern of evolution for the phylogenetic niche conservatism relating to aforementioned traits (46, 64). Accordantly, many other studies had shown that robust phylogenetic signal was related to different variables within short distances across ecosystems of diverse types (47, 50, 65–67). On this basis, recent advanced quantitative ecological models (47, 51, 68), can be applied to compare observed between-community diversity of both phylogeny and taxonomy against a null-model, thereby quantify the relative importance of ecological processes underlying the gut microbiota in response to feed-borne bacteria.

The turnover of an ecological community (fugu gut microbiota in our case) is drived by two forces, determinacy and stochasticity (69). Determinacy indicates that the presence/absence and relative abundances of species is governed by abiotic and biotic factors, which are associated with ecological selection (niche effect) (70). In contrast, stochasticity indicates that the relative abundances (ecological drift) of species are determiend by probabilistic dispersal and randomness (neutral effect), which are not caused by environmentally determined fitness (69, 71). Our Mantel’s test results shown that severe differences in feed source signals act as a strong habitat filter and lead to significant phylogenetic clustering (|βNTI| >2), whereas the degree to which community composition influenced by such filtering was weakened under moderate feed source signal (Figure 4A). This result further emphasized that niche effect posed a strong selection pressure in determining the community assembly (71), thereby driving bacterial communities into a phylogenetically more closed cluster (64). As a continual inoculum, feeding poses a significant destabilizing force in fish intestinal microbiota and results in a continuous community turnover (18, 19). On this basis, community invasibility, the successful colonization of allochthonous organisms such as feed-borne bacteria in a given community like fish gut, can be indicative of community stability (72, 73). In supporting this notion, we found that with the changes in feed-signals increases, the assembly processes of gut bacterial community were gradually shifted from homogeneous selection, to stochasticity, then to variable selection (Figure 4A). Of note, such pattern was more obviously observed for rare sub-community (Mantel R=0.492, Figure 4C) than to abundant one (Mantel R=0.275, Figure 4B), indicating the abundant bacterial taxa (mostly *gamma-proteobacterial* copiotroph) might be more competitively dominant and persistent in gut niche. Specifically, the variable selection process could be possibly explained by the increased habitat complexities of tripartite interactions among host, bacteria and nutrient, considering accumulated ingested feed-borne bacteria can cause more disturbance, thereby imposed a larger impact on rare sub-community as was evident by the increased alpha-diversity in slow growth fugu (more prone to feed disturbance). As for the homogeneous selection, we infered that the deterministic factors, particularly by the host and autochthonous bacteria became more dominant and stable (e.g. lower alpha-diversity in fast growth fugu) as the changes of feed source signal reduced, thereby jointly exerted more convergent force of selection.

By inferring the turnover in ESV composition (RCbc), the relative importance of stochastic processes can be further fractionated into homogenizing dispersal, dispersal limitation, and the undominated fraction (74). We observed a large fraction of community shift was governed by dispersal limitation (~40%), whereas very little (3%) by homogenizing dispersal (Figure 4D). Higher level of dispersal limitation can be viewed as a result of host effect, by which the degree to which bacteria move among individual gut communities was limited, even though the influence from the feed is strong. In line with these, both the fitness of neutral model (R2=0.154) and migration rate (m=0.012) for abundant sub-community were lower than that of rare sub-community (R2=0.639; m=0.057), further emphasizing the limited levels of dispersal leads to dissipation of diversity into local gut communities, thus decreased alpha diversity (71). The lower degree of homogenizing dispersal (75) is consistent with our observed higher degree of beta dispersion in fugu gut microbiota. In addtion, an undominated fraction was also observed for all sub-communities, particlularly to a larger extent in abundant one (49%). The inflated undominated fraction could be viewed as the results of diminished selection and/or dispersal rates, that both are opposite to that of variable selection (74). As such, it is tempting to speculate that the observed undominated fraction here is likely reflecting the inconsistent selection from either feed-borne bacteria and host filtering (or both), which possibly due to the varied host physiology (e.g. gastric evacuation and growth) and/or gut nutrition (e.g. feeding fluctuation and frequency) (19, 59, 76).

In conclusion, this snapshot study shown here characterized the compositional, functional and phylogenetical stability of gut microbiota from farmed fugu within a local pond environment. This stability is accompanied largely by homogeneous selection and dispersal limitation, thereby, a reduced intraspecies competition, enabling community stability and partial variations among hosts. However, presumptive attaching bacteria introduced by feed can pose a strong restructuring force upon gut bacterial communities. Such disturbance involved with variable selection, therefore, an augmented interaction between autochthonous and allochthonous species, entailing community unstability and shift. Moreover, we observed marked alterations in the composition and function of gut microbiota in slow growth host, potentially correlated to the different host physiological conditions, e.g. gastric evacuation rate and intestinal transit time of digesta. Nevertherless, we can not completely rule out the possiblity that such community shift in gut microbiota can be due to the efflux DNA of feed borne bacteria, considering the lack of measurement of active bacteria for both gut microbiota and feed in our experiment. Future longitudinal studies concerning the ingested feed-borne bacteria and its interactions with both gut symbiosis and its host are needed to determine whether the observed correlations can be maintained or any causality can be confirmed. Altogether, our findings emphasized the intricate linkage between feed and gut microbiota, and highlighted the essential prerequisite to resovle the signal from feed-associated bacteria before the conclusions of comparative analysis of microbiota can be drawn.

## 4. MATERIALS AND METHODS

### 4.1. Sample collection

We collected samples from 7-month-old juvenile obscure puffer (*T. obscurus*) in December 2017 from a single commercial supplier and raised in a fishfarm located in Yangzhong, East China. After acclimating to commercial pellet feeds, the fish offsprings from the full-sib family were introduced into indoor greenhouse ponds (80m x 25m x 1.8m) with a stocking density of 25 thounds per pond since April. Each pond had a separate water inlet and outlet. Three adjacent ponds were selected as the objects of study. The practical pond management is identical in terms of daily feeding rate (twice per day, i.e. 10am and 2pm), and pond water exchange rate (twice per month, 10%~20%). Approximately 20 post-feeding (1 day) fishes per pond were randomly captured using a fishing net along the different sites of each pond. No obvious fin damage or skin injury were observed across the body among indivisuals. Fishes were anesthetized with 0.05% MS-222 (Sigma-Aldrich, St. Louis, MO, USA), and immediatedly weighted and fork length measured. After being surface sterilized with 75% ethanal, the 2 cm long part of distal intestine was dissected out to collect the gut content using sterilized spatula. Subsequently, other tissues including the whole liver and gonad were also dissected and rapidly weighted. All samples were placed in individually labelled tubes and kept well in dry ice before shipping. All animal experiments were performed following the protocol approved by the Ethics Committee of Experimental Animals at Hohai University, and were in direct accordance with the Animal Care Guidelines issued by the Ministry of Science and Technology, the People’s Republic of China.

For the environmental samples including pond water and sediment, three longitudinal distributed sites (similar locations among the ponds) per pond were chose respectively. Approximately 2.0 L of pond water was collected from 10 to 20 cm below surface per sites within each pond respectively. Water sample was prefilted by 100-μm pore sized nylon mesh, and the flowthrough was further filtered with 0.22um MilliPore membrane with vacuum pump to collected the water planktonic microbes. The sediment was collected from the same site as water sample, and was packaged in airtight sterile plastic bags. All the microbial samples were preseved with dry ice during the shipment and stored at −80°C before use. The detailed metadata for each sample was recored in Table S3.

### 4.2 DNA isolation and bacterial 16S rRNA amplicon sequencing

All the fugu gut contents (n = 61), feed pellet (n=3), water (n=9) and sediment (n=9) samples were subjected to bacterial 16S rRNA amplicon profiling by Illumina sequencing. DNA was extracted using FastDNA™ SPIN Kit for Soil (MP Biomedicals, Irvine, CA, USA) according to manufacturer’s instructions, and was verified by 0.8% agarose gel electrophoresis. The integrity and concentration of DNA were determined by Nanodrop ND 2000 spectrophotometer (Thermo Fisher Scientific Inc., Waltham, MA, USA) and Quant-iT PicoGreen dsDNA assay kit (Thermo Fisher Scientific Inc., Waltham, MA, USA), respectively. The V4-V5 hypervariable region of bacterial16S rRNA gene were amplified using the primer pair 515F (5’-CTGCCAGCMGCCGCGGTAA-3’) and 926R (5’-CCGTCAATTCMTTTRAGTTT-3’), and the unique eight-nucleotide barcode sequences were incorporated into each sample. Each sample was tested in triplicate (together with No template controls) with the 25-μl reaction mix consisted of 40 ng template DNA, 5 μl reaction buffer (5x), 5 μl GC buffer (5x), 2 μl dNTP (2.5mM), 10 pM of barcoded forward and reverse primers, and 0.75 U Q5^®^ High-Fidelity DNA Polymerase (NEB, Ipswich, MA, USA). The thermocycling program consisted of one hold at 98 °C for 2 min, followed by three-step 25 cycles of 15 s at 95 °C, 30 s at 55 °C and 30 s at 72 °C, and one final fold at 72°C for 5 min.

All PCR amplicons were subjected to 2% agarose gel electrophoresis, and purified with the AMPure XP Kit (Beckman Coulter GmbH, USA), quality checked using Agilent High Sensitivity DNA Kit (Agilent, Santa Carla, CA, USA) and Quant-iT PicoGreen dsDNA assay kit. DNA libraries for purified PCR amplicons were constructed using an Illumina TruSeq Nano DNA LT Library Prep Kit (Illumina, San Diego, CA, USA) according to the manufacturer’s instruction, and 300-bp pair-ended insertion were sequenced using MiSeq Reagent Kit V3 chemistry on Illumina MiSeq platform owned by Personal Biotechnology Co., Ltd. (Shanghai, China).

### 4.3 Reads processing and bacterial 16S rRNA gene profiling

The adapters (indexers) in the paired-end raw reads were trimmed out by the quality control tool Trim-Galore (a wrapper tool based on Cutadapt v.1.4.2 and FastQC v.0.10.1) for high throughput sequence data, as set up by default quality threshold of Q20 (77). The FastQC reports from both before- and after-trimming were checked. Further, the 16S rRNA amplicon raw reads were processed according to the previous literatures (78, 79) using USEARCH v.10.0 (80). In brief, the paired-end reads were merged and relabeled using ‘-fastq_mergepairs’; both barcodes and primers were removed using ‘-fastx_truncate’; reads with low-quality (error rates >0.01) and redundancy were filtered and dereplicated using ‘-fastq_filter’ and ‘-fastx_uniques’, respectively. The *de novo* biological sequences, i.e. ESVs (exact sequence variants) were clustered and chimeras were filtered using ‘-unoise3’. Subsequently, the operational taxonomic units (OTUs) table was created by ‘-otutab’. The taxonomy for each representative sequences were assigned by the sintax algorithm using the Ribosomal Database Project (RDP) classifier (RDP training set v16) (81, 82), and chloroplast ESVs were removed after taxonomy-based filtering. Sequencing depth was normalized by subsample using ‘otutab_norm’, yielding 7,000 sequences per sample to keep all the samples for downstream analysis. Community diversity was analyzed using ‘-alpha_div’, ‘-alpha_div_rare’, ‘-cluster_agg’ and ‘-beta_div’ respectively.

### 4.4 Featured taxa identifying and source tracking across different bacterial communities

To discriminate the compositional difference and identify the featured taxa across different bacterial communities, R package RandomForest (v.4.6-14)(83) was used to classify the relative abundances of bacterial taxa in the family level across different sample source using parameters of ‘ntree = 1000, importance = TRUE, proximity = TRUE’, followed by cross-validation using *rfcv()* function for feature selection. Important features were clustered according to the NCBI taxonomy and visualized by Interactive Tree of Life, iTOL (84).

To further identify the potential origins for the compositional structure for each indicated sample group, SourceTracker (85) was used to estimate the proportions of source bacteria from environments in each community. All the samples (n=82) were first rarefied at 1000 sequences as training set, and SourceTracker object was trained by *sourcetracker()* function. Each indicated sample group was used as test set and of which the source proportions (from SourceTracker object) were estimate by *predict()* function with the alpha values tuned to 0.001.

The ESV table was further analyzed to identify featured bacterial taxa that are specific to indicated group. ESVs were filtered with abundance greater than 0.0001% across gut samples. Significant changes in relative ESV abundance were identified with threshold of logarithmic LDA (linear discriminant analysis) score greater than 2 combining with effect size (LEfSe) algorithm (86).

### 4.5 Phenotypic infering of bacterial communities

For metagenomics inference, the ESVs table normalized by 16S rRNA copy numbers were annotated by PICRUSt (87) with closed reference picking (Greengenes v.13_5)(88). Abundance of bacterial communities with two trophic modes of bacterial life namely copiotroph and oligotroph were calculated by computing the mean number of ribosomal operon copies in the genome across all bacterial ESVs presented in each sample as previously described (89, 90). Firstly, rRNA copy number per ESVs (predicted by PICRUSt) within a given sample was multiplied by the corresponding relative abundance and summed for the total copy number per sample. Secondly, mean community copy number was averaged per indicated sample group (fish gut and environments) and were subjected to group-wised multiple comparisons using indicated statistical approach as will be described below. Biologically interpretable phenotypes such as oxygen tolerance, Gram staining and pathogenic potential, within each indicated community were predicted based on the Greengenes OTU table using BugBase (91). Taxonomic level was set as bacterial family.

### 4.6 Quantifying relative importance of ecological processes in gut bacterial community assemblage

The existence and degree of phylogenetic relatedness in species traits were assessed through the calculation of the phylogenetic signal of a given trait. To this end, both Blomberg’s K statistic (46) and Mantel’s Correlogram (45) were used to measure phylogenetic signal using *phyloSignal()* and *phyloCorrelogram()* implemented in R package ‘Phylosignal’ (92). To assess the phylogenetic structure of gut bacterial communities, the standardized effect size of mean nearest taxon distances (ses.MNTD) per community was calculated using *ses.mntd()* function implemented in ‘Picante’ package (93), supplied with options of “taxa.labels” as null model, unweighted and 999 randomization. Ses.MNTD values that were significant less or greater than 0 indicate phylogenetically close-related or unrelated than expected by chance, respectively (64). Ecological null model was used to quantify the relative importance of ecological processes in driving microbial community assemblage (51). Briefly, both metrics of β-nearest taxon index (βNTI) and Bray-Curtis-based Raup-Crick (RC_BC_) were used to quantify the contributions from deterministic and stochastic processes, respectively. First, the phylogenetic beta diversity were measured by computing unweighted inter-community MNTD metric (βMNTD) using the *comdistnt()*. Further, the βNTI, i.e. the degree to which observed βMNTD deviates from the mean of the null distribution (999 randomization) was computed according to Stegen et al. (51). The influence of deterministic community turnover including homogeneous selection and variable selection (70) is estimated by infering the fraction of pairwise community comparisons with significance thresholds of βNTI < −2 and βNTI > 2, respectively (51). In turn, the non-significant fractions with |βNTI|< 2 that indicate community turnover governed by stochastic ecological processes, were further subdivided according to the taxonomic β-diversity metrics of RC_BC_. Within this subset (|βNTI|< 2), RC_BC_ metric normalizes the deviation between observed Bray–Curtis and the null distribution (9,999 randomization) to vary between −1 and +1. The turnover of microbial community with significant RC_BC_ values that are less than −0.95 or greater than +0.95 were interpreted as governed by homogenizing dispersal and dispersal limitation, respectively. The ones with non-significant values (i.e. |RC_BC_| 0.95) was regarded as community turnover governed by undominated processes without a profound effect by any given ecological pressure (i.e., weak dispersal and weak selection).

### 4.7 Statistics

All statistical analyses were performed under R environment (v3.6.0) through RStudio (v1.1.463). Before analysis, the normality and homogeneity of variances of datasets were tested using *shapiro.test()* and *bartlett.test()*, respectively. In turn, parametric or non-parametric tests were applied accordingly. For parametric datasets, group means were compared by one-way ANOVA using *aov()*, followed by *post-hoc* test using *TukeyHSD()*. For non-parametric datasets, rank sum were compared by Kruskal-Wallis test using *kruskal.test()*, followed by Wilcoxon rank sum test using *wilcox_test()*. Significance were considered while FDR-adjusted *P* values were less than 0.05, and the detailed statistical test applied in each dataset is elaborated in the figure legend.

The homogeneity of multivariate dispersion of groups was evaluated by (PERMDISP) test with *betadisper()* followed by permutation test with *permutest()*. Ordination (Principle Coordinate analyses, PCoA) were performed based on Bray-Curtis dissimilarity metric using *cmdscale()* followed by permutational multivariate analyses of variance (PERMANOVA) test by *adonis()* inplemented in ‘vegen’ package (94). The clustering analysis based on individual fish body mass were determined by *eclust ()* using K-means distance and *clusGap()* conducting gap statistic algorithm implemented in ‘factoextra’ package (95), followed by visualization according to principal components analysis by *prcomp()*. Spearman-correlations between the phylogenetic beta diversity (βNTI matrix) and indicated traits difference (euclidean-based trait distance martix) we assessed using *mantel()* implemented in ‘ecodist’ package (96) with 9,999 permutations. The *neutral.fit()* in the ‘MicEco’ package (97) was used to evaluate the fitness of neutral model (98) on the gut bacterial community, and to determine the contribution of neutral processes in community assembly.

## AVALIABILITY OF DATA

The data set generated and analyzed for this study is available in the NCBI sequencing reads archive (SRA), under BioProject accession number PRJNA658467.

## AUTHOR CONTRIBUTIONS

XJ conceived and designed the experiments. ZZ, YS provided the initial framework and aquired fund for the study. XJ conducted the sampling and experimental work. XJ and ZC performed data analysis and visualization. XJ, ZC, YS, JFG and ZZ interpreted the results. XJ wrote the manuscript with input of all authors. All authors read and approved the final manuscript.

## ACKNOWLEDGEMENTS

This work was supported supported by the National Key Research and Development Program of China (2018YFD0900200); the National Natural Science Foundation of China (31872597); the Fundamental Research Funds for the Central Universities from China (B200202141); and the Earmarked Fund for Jiangsu Agricultural Industry Technology System (JATS [2019]477). We thank Mr Zhu Jikun for his deep knowledge of pufferfish husbandry, and Song Jing, Gao Tianheng, Gao Fanxiang, Wu Chao, Liu Yupeng and Wen Duan for their help in sampling.

We declare that we have no competing interests.

## SUPPLEMENTAL MATERIAL

### TABLE LEGENDS

**TABLE.S1.** Numbers of reads that had passed quality control for each sample.

| ID | Size | ID | Size | ID | Size | ID | Size |
| --- | --- | --- | --- | --- | --- | --- | --- |
| F1 | 36736 | F9 | 26043 | Fb9 | 25070 | FD1 | 36815 |
| F10 | 22333 | Fb1 | 25583 | Fc1 | 21975 | FD2 | 37651 |
| F11 | 27447 | Fb10 | 24496 | Fc10 | 62291 | FD3 | 38229 |
| F12 | 28627 | Fb11 | 23487 | Fc11 | 24664 | S1In | 10336 |
| F13 | 35318 | Fb12 | 22294 | Fc12 | 22261 | S1Mid | 9619 |
| F14 | 33645 | Fb13 | 27105 | Fc13 | 55646 | S1Out | 9122 |
| F15 | 31020 | Fb14 | 20734 | Fc14 | 28143 | S2In | 10294 |
| F16 | 32521 | Fb15 | 23462 | Fc15 | 24754 | S2Mid | 8410 |
| F17 | 31784 | Fb16 | 25183 | Fc16 | 24317 | S2Out | 12520 |
| F18 | 30501 | Fb17 | 27676 | Fc17 | 26723 | S3In | 8888 |
| F19 | 33761 | Fb18 | 26288 | Fc18 | 23858 | S3Mid | 9550 |
| F2 | 30753 | Fb19 | 24460 | Fc19 | 27784 | S3Out | 7753 |
| F20 | 25821 | Fb20 | 24798 | Fc2 | 23387 | W1In | 47616 |
| F21 | 26256 | Fb21 | 26507 | Fc20 | 59896 | W1Mid | 25924 |
| F3 | 31846 | Fb22 | 21946 | Fc3 | 21910 | W1Out | 22123 |
| F4 | 28077 | Fb3 | 23116 | Fc4 | 28096 | W2In | 16548 |
| F5 | 29201 | Fb5 | 29513 | Fc5 | 27971 | W2Mid | 23136 |
| F6 | 30302 | Fb6 | 26927 | Fc6 | 20612 | W2Out | 30361 |
| F7 | 28165 | Fb7 | 31320 | Fc7 | 23442 | W3In | 33120 |
| F8 | 36612 | Fb8 | 22443 | Fc8 | 22271 | W3Mid | 28363 |
|  |  |  |  | Fc9 | 66205 | W3Out | 29713 |

**TABLE.S2.** Statistical results for each indicated figure.

**Figure 1A**
| **(1) adonis setting pond as factor: adonis (dis ~ Pond, data = design, by = NULL)** |  |  |  |  |  |  |
| --- | --- | --- | --- | --- | --- | --- |
|  | Df | SumsOfSqs | MeanSqs | F.Model | R2 | Pr(>F) |
| Pond | 2 | 2.9104 | 0.97014 | 3.2346 | **0.11064** | **0.001** |
| Residuals | 79 | 23.3941 | 0.29992 | 0.88936 |  |  |
| Total | 81 | 26.3045 | 1 |  |  |  |
| **(2) adonis setting sample groups (F, Fd, W, S) as factor: adonis (dis ~ group, strata = design\$Pond, data = design, by=NULL)** | | | | | | |
|  | Df | SumsOfSqs | MeanSqs | F.Model | R2 | Pr(>F) |
| group | 3 | 6.2479074 | 2.0826358 | 8.09934898 | **0.2375221** | **0.001** |
| Residuals | 78 | 20.0566234 | 0.2571362 | NA | 0.7624779 | NA |
| Total | 81 | 26.3045308 | NA | NA | 1 | NA |

**Figure 1B**
| Pairwise comparisons using Wilcoxon rank sum test, P value adjustment method: BH |  |  |  |  |  |  |  |  |  |
| --- | --- | --- | --- | --- | --- | --- | --- | --- | --- |
|  | F.F | F.Fd | F.S | F.W | Fd.Fd | Fd.S | Fd.W | S.S | W.S |
| F.Fd | 0.00010227 | NA | NA | NA | NA | NA | NA | NA | NA |
| F.S | 1.32E-245 | 7.18E-81 | NA | NA | NA | NA | NA | NA | NA |
| F.W | 1.72E-249 | 7.25E-82 | 1.16E-05 | NA | NA | NA | NA | NA | NA |
| Fd.Fd | 0.00330435 | 0.00342397 | 0.00329105 | 0.00329105 | NA | NA | NA | NA | NA |
| Fd.S | 2.25E-16 | 1.34E-14 | 3.66E-06 | 9.11E-05 | 0.00186231 | NA | NA | NA | NA |
| Fd.W | 1.12E-15 | 1.34E-14 | 5.41E-09 | 1.44E-06 | 0.00421957 | 1.54E-10 | NA | NA | NA |
| S.S | 2.90E-13 | 2.18E-12 | 3.40E-23 | 3.40E-23 | 0.00502153 | 1.70E-11 | 2.28E-11 | NA | NA |
| W.S | 2.08E-11 | 0.28852238 | 4.18E-45 | 4.75E-47 | 0.00356081 | 1.53E-14 | 1.61E-14 | 2.17E-17 | NA |
| W.W | 1.41E-10 | 1.08E-09 | 3.40E-23 | 3.40E-23 | 0.00056274 | 1.70E-11 | 2.28E-11 | 0.16972718 | 2.17E-17 |

**Figure2A**
| The quality of representation (cos2) for each indicated body mass indice on the principal components (PC1~4) |  |  |  |  |
| --- | --- | --- | --- | --- |
|  | PC1 | PC2 | PC3 | PC4 |
| Weight.g | 0.962038403 | 0.025239425 | 0.003224548 | 0.009497624 |
| Length.cm | 0.917584928 | 0.012917832 | 0.068747184 | 0.000750056 |
| Weigh.gonad.g | 0.718414987 | 0.280904081 | 0.000676607 | 4.33E-06 |
| Weigh.liver.g | 0.925801286 | 0.036780732 | 0.032483293 | 0.004934688 |

**Figure 2B**
| (1) adonis analysis on feed proportions predicted by SourceTracker as continuous variable, and pond as strata: adonis (dis ~ feed, strata = design\$Pond, data = design, by = NULL) | | | | | | |
| --- | --- | --- | --- | --- | --- | --- |
|  | Df | SumsOfSqs | MeanSqs | F.Model | R2 | Pr(>F) |
| feed | 1 | 6.67712586 | 6.67712586 | 50.4357901 | 0.46087107 | 0.001 |
| Residuals | 59 | 7.81093 | 0.13238864 | NA | 0.53912893 | NA |
| Total | 60 | 14.4880559 | NA | NA | 1 | NA |
| (2) adonis analysis on body mass index (PC1) as continuous variable, and pond as strata: adonis (dis ~ Comp.1, strata = design\$Pond, data = design, by = NULL) | | | | | | |
|  | Df | SumsOfSqs | MeanSqs | F.Model | R2 | Pr(>F) |
| Comp.1 | 1 | 0.97960187 | 0.97960187 | 4.27854366 | 0.06761445 | 0.003 |
| Residuals | 59 | 13.508454 | 0.22895685 | NA | 0.93238555 | NA |
| Total | 60 | 14.4880559 | NA | NA | 1 | NA |

**Figure 2C**
| Multiple comparisons of proportions of feed (predicted by SourceTracker) using TukeyHSD. |  |  |  |  |
| --- | --- | --- | --- | --- |
|  | diff | lwr | upr | p adj |
| Medium-Fast | 0.106935 | -0.1083305 | 0.32220053 | 0.461004 |
| Slow-Fast | 0.22879706 | 0.00340915 | 0.45418497 | 0.04588741 |
| Slow-Medium | 0.12186206 | -0.1126851 | 0.35640922 | 0.4292129 |

**Figure 2D**
| Pairwise comparisons using Wilcoxon rank sum test (P value adjustment method: BH ) |  |  |  |  |  |  |  |  |
| --- | --- | --- | --- | --- | --- | --- | --- | --- |
|  | Fst-Fd | Fst-Fst | Fst-Med | Fst-Slw | Med-Fd | Med-Med | Med-Slw | Slw-Fd |
| Fst-Fst | 1.05E-05 | NA | NA | NA | NA | NA | NA | NA |
| Fst-Med | 7.01E-07 | 0.74256729 | NA | NA | NA | NA | NA | NA |
| Fst-Slw | 0.00036736 | 0.07494946 | 0.01044206 | NA | NA | NA | NA | NA |
| Med-Fd | 0.11633715 | 0.09130052 | 0.03107713 | 0.35913902 | NA | NA | NA | NA |
| Med-Med | 1.05E-06 | 0.69882918 | 0.86343745 | 0.03107713 | 0.03107713 | NA | NA | NA |
| Med-Slw | 3.08E-06 | 0.69882918 | 0.36686535 | 0.14483626 | 0.0693612 | 0.36686535 | NA | NA |
| Slw-Fd | 0.00036736 | 0.22086864 | 0.36686535 | 0.06239833 | 0.09560009 | 0.37318151 | 0.28345795 | NA |
| Slw-Slw | 1.33E-05 | 0.92290083 | 0.86343745 | 0.12770101 | 0.07494946 | 0.77457806 | 0.69882918 | 0.36686535 |

**Figure 2E**
| Pairwise comparisons using Wilcoxon rank sum test (P value adjustment method: BH ) |  |  |  |  |  |
| --- | --- | --- | --- | --- | --- |
|  | Fd | Fst | Med | Sdm | Slw |
| Fst | 0.011500301 | NA | NA | NA | NA |
| Med | 0.011779595 | 0.804463546 | NA | NA | NA |
| Sdm | 0.022131884 | 9.18E-05 | 9.18E-05 | NA | NA |
| Slw | 0.02484192 | 0.04352426 | 0.072248752 | 0.000105016 | NA |
| Wtr | 0.022131884 | 9.18E-05 | 9.18E-05 | 0.000877415 | 0.000105016 |

**Figure 4 Blomberg's K statistical output.**
| #ALL_fugu | K | PIC.variance.obs | PIC.variance.rnd.mean | PIC.variance.P | PIC.variance.Z |
| --- | --- | --- | --- | --- | --- |
| PC1 | 0.05398214 | 61.668551 | 205.5198 | 0.001 | -8.668789 |
| feed | 0.05049687 | 2.065222 | 6.30681 | 0.001 | -8.897068 |
| #Fast_fugu | K | PIC.variance.obs | PIC.variance.rnd.mean | PIC.variance.P | PIC.variance.Z |
| PC1 | 0.04644161 | 16.148637 | 47.220684 | 0.001 | -5.047588 |
| feed | 0.08689312 | 1.788734 | 9.770652 | 0.001 | -8.297465 |
| #medium_fugu | K | PIC.variance.obs | PIC.variance.rnd.mean | PIC.variance.P | PIC.variance.Z |
| PC1 | 0.02844361 | 9.942098 | 15.908873 | 0.001 | -3.913017 |
| feed | 0.05284201 | 3.271343 | 9.773969 | 0.001 | -7.303169 |
| #slow_fugu | K | PIC.variance.obs | PIC.variance.rnd.mean | PIC.variance.P | PIC.variance.Z |
| PC1 | 0.03317783 | 5.482231 | 11.00622 | 0.001 | -3.705141 |
| feed | 0.05582301 | 1.262326 | 4.26296 | 0.001 | -8.018897 |

**TABLE.S3.**
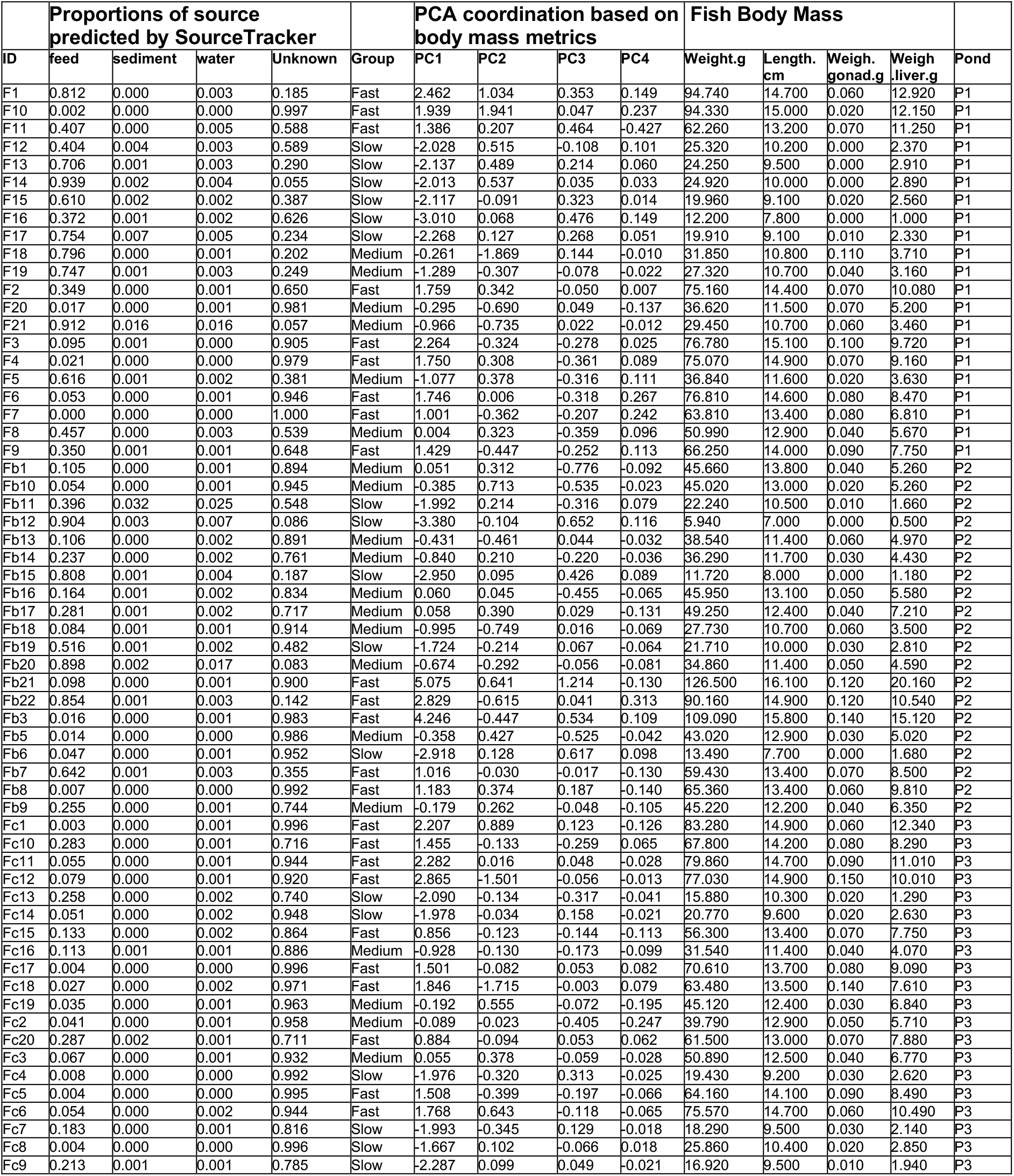
Metadata.

**TABLE.S4.** Importance value of Random-Forest predicted bacterial families in classfying sample groups.

| Family | F | FD | S | W |
| --- | --- | --- | --- | --- |
| Cytophagaceae | -0.512 | -0.512 | -0.475 | 1.500 |
| Chitinophagaceae | -0.792 | -0.782 | 1.298 | 0.275 |
| Conexibacteraceae | -0.768 | -0.772 | 0.209 | 1.331 |
| Caulobacteraceae | -0.814 | -0.794 | 0.349 | 1.259 |
| **Mycoplasmataceae** | 1.500 | -0.499 | -0.500 | -0.500 |
| Oxalobacteraceae | -0.859 | -0.866 | 0.978 | 0.747 |
| Comamonadaceae | -0.777 | -0.781 | 0.243 | 1.314 |
| Cryomorphaceae | -0.609 | -0.610 | -0.260 | 1.479 |
| **Brevinemataceae** | 1.500 | -0.500 | -0.500 | -0.500 |
| Methylocystaceae | -0.812 | -0.818 | 0.401 | 1.229 |
| Sphingomonadaceae | -0.681 | -0.680 | -0.077 | 1.438 |
| Mycobacteriaceae | -0.521 | -0.521 | -0.458 | 1.499 |
| Planctomycetaceae | -0.542 | -0.555 | 1.496 | -0.399 |
| Rhodobacteraceae | -0.740 | -0.843 | 1.286 | 0.297 |
| Rhodocyclaceae | -0.507 | -0.508 | 1.500 | -0.485 |
| Saprospiraceae | -0.841 | -0.838 | 1.139 | 0.540 |
| Acidimicrobiaceae | -0.683 | -0.739 | 1.410 | 0.012 |
| Chromatiaceae | -0.793 | -0.793 | 1.284 | 0.302 |
| Opitutaceae | -0.622 | -0.622 | 1.474 | -0.231 |
| (Unassigned) | -0.879 | -0.714 | 1.270 | 0.323 |
| Flavobacteriaceae | -0.691 | -0.705 | -0.027 | 1.423 |
| Verrucomicrobiaceae | -0.662 | -0.659 | -0.132 | 1.453 |
| **Rikenellaceae** | 1.500 | -0.500 | -0.500 | -0.500 |
| Burkholderiaceae | -0.580 | -0.260 | -0.640 | 1.479 |
| Geobacteraceae | -0.518 | -0.463 | 1.499 | -0.518 |
| Microbacteriaceae | -0.509 | -0.517 | -0.474 | 1.500 |
| Xanthomonadaceae | -0.668 | -0.681 | 1.443 | -0.094 |
| Methylococcaceae | -0.596 | -0.600 | 1.484 | -0.287 |
| Planococcaceae | -0.490 | 1.500 | -0.504 | -0.506 |
| Carnobacteriaceae | -0.500 | 1.500 | -0.503 | -0.497 |
| Idiomarinaceae | -0.500 | 1.500 | -0.500 | -0.500 |
| Bacillaceae_2 | -0.500 | 1.500 | -0.500 | -0.500 |
| Oceanospirillaceae | -0.500 | 1.500 | -0.500 | -0.500 |
| Acidaminococcaceae | -0.529 | 1.498 | -0.547 | -0.423 |

**TABLE.S5.** Mean relative abundance of bacterial classes from each indicated sample group.

| Order | Fst | Med | Slw | Fd | Wtr | Sdm |
| --- | --- | --- | --- | --- | --- | --- |
| **Gammaproteobacteria** | 43.4 | 42.2 | 61.8 | 80.4 | 7.8 | 15.3 |
| Spirochaetia | 17.8 | 18.9 | 11.2 | 0.0 | 0.0 | 0.4 |
| Bacteroidia | 16.5 | 15.8 | 8.6 | 0.5 | 0.0 | 1.3 |
| Mollicutes | 9.9 | 12.0 | 5.7 | 0.0 | 0.0 | 0.0 |
| Betaproteobacteria | 0.3 | 0.4 | 0.6 | 1.1 | 19.8 | 21.5 |
| Actinobacteria | 0.1 | 0.2 | 0.5 | 0.2 | 33.6 | 3.5 |
| Clostridia | 4.8 | 3.4 | 3.8 | 3.4 | 0.0 | 0.4 |
| Unassigned | 2.2 | 2.7 | 2.4 | 1.6 | 0.7 | 10.5 |
| Alphaproteobacteria | 0.1 | 0.2 | 1.3 | 0.3 | 13.8 | 9.7 |
| Cytophagia | 0.0 | 0.0 | 0.1 | 0.0 | 10.9 | 3.0 |
| Bacilli | 1.0 | 1.7 | 1.5 | 12.2 | 0.2 | 0.0 |
| Other | 4.0 | 2.5 | 2.5 | 0.3 | 13.2 | 34.5 |

**TABLE.S6.**
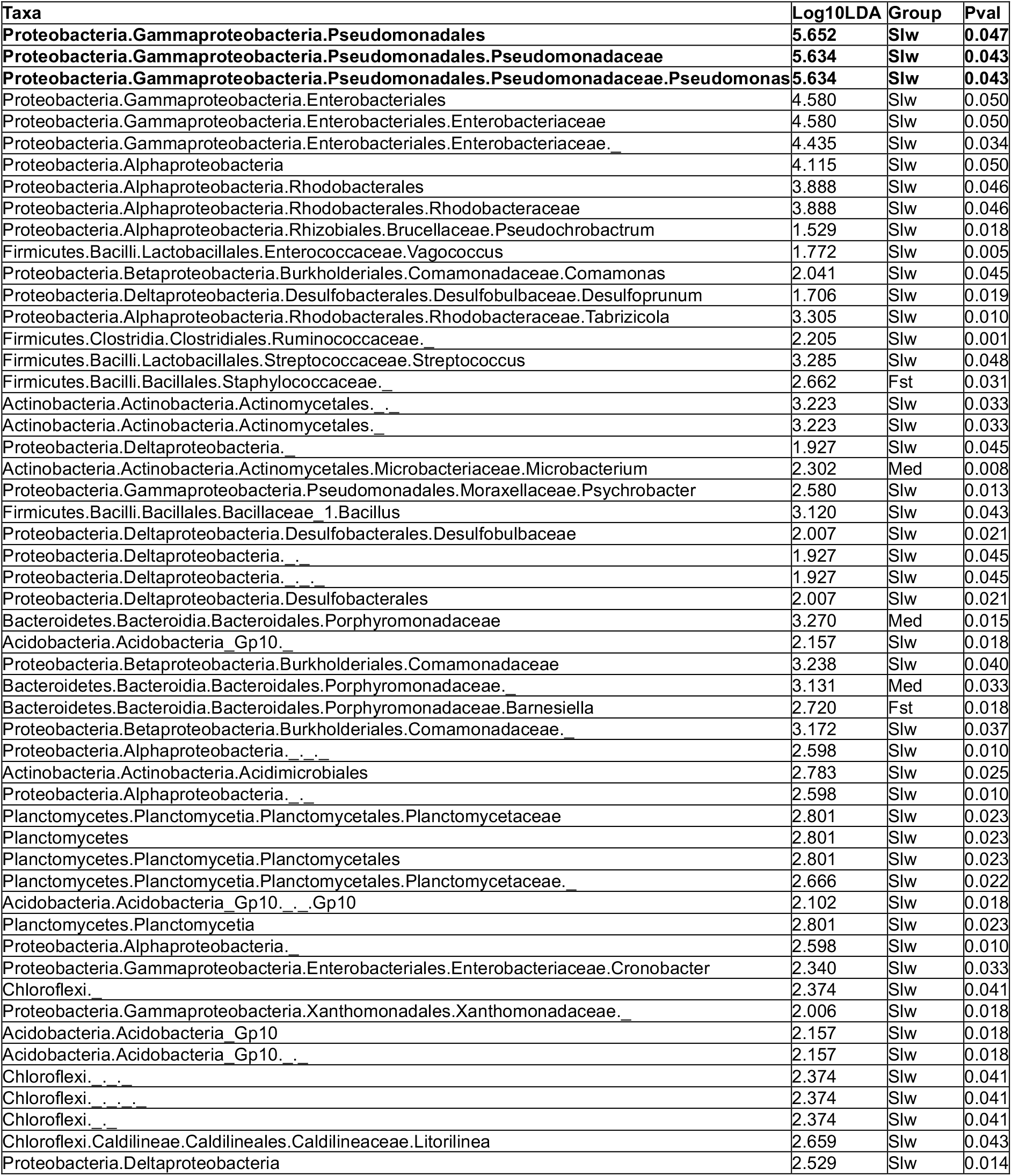
Discriminative bacterial taxa identified by LEfSe.

### FIGURE LEGENDS

**Figure S1.**
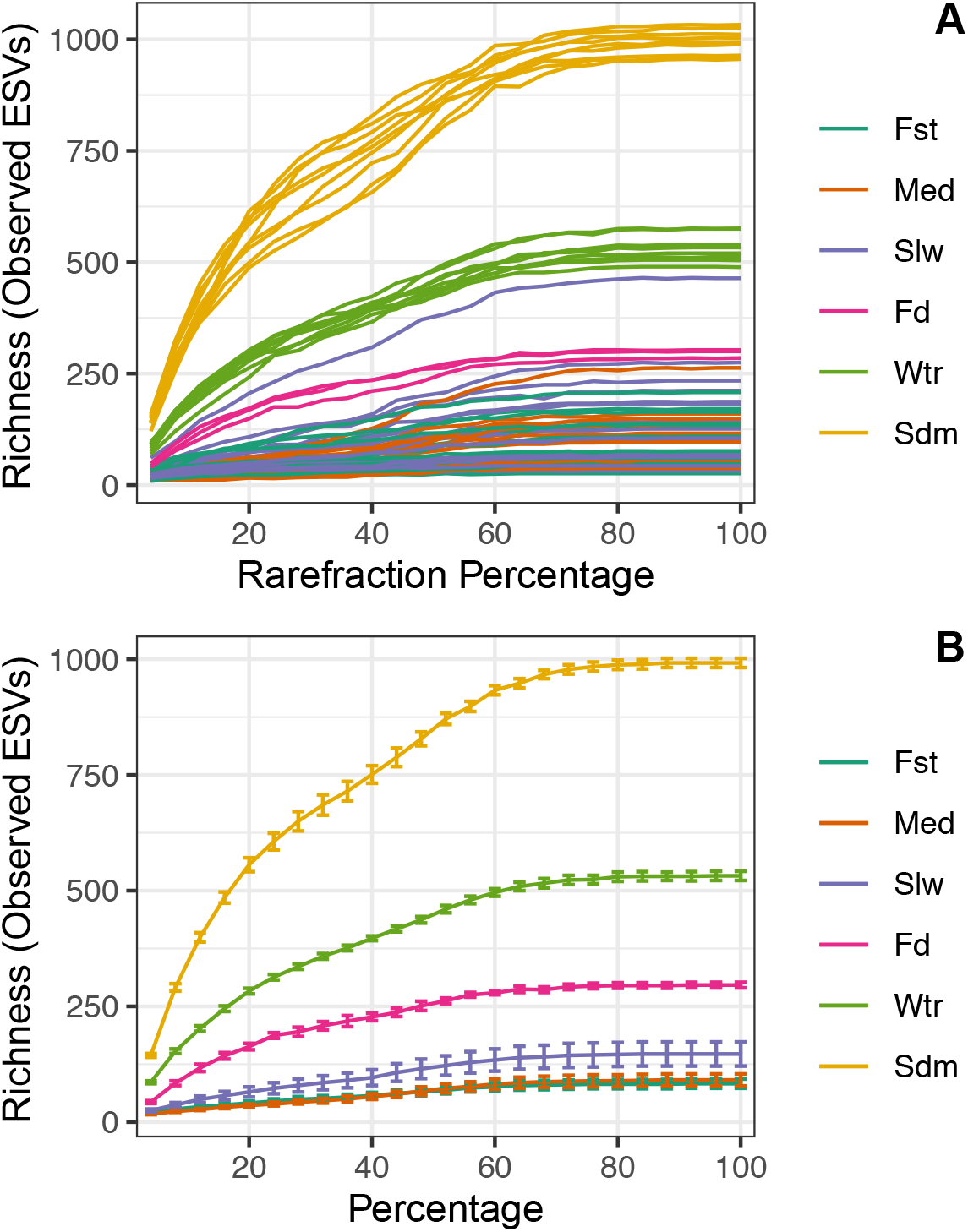
Evalution of sequencing depth of bacterial communities across different sample groups. **(A)** Rarefaction curves of detected bacterial species (numbers of observed ESVs) within each indicated microbiota sample tend to be ‘flat’ as the numbers of sampling increase, suggesting the sampling covers the most bacterial species (saturated). **(B)** Rarefaction curves of detected bacterial species (numbers of observed ESVs) in each indicated sample groups. Vertical bars denote standard error. Samples size for each indicated groups are as follows: fish (n=61), including fast ‘Fst’ (n=24), Medium ‘Med’ (n=20) and Slow ‘Slw’ (n=17); feed ‘Fd’ (n=3); water ‘Wtr’ (n=9); sediment ‘Sdm’ (n=9).

**Figure S2.**
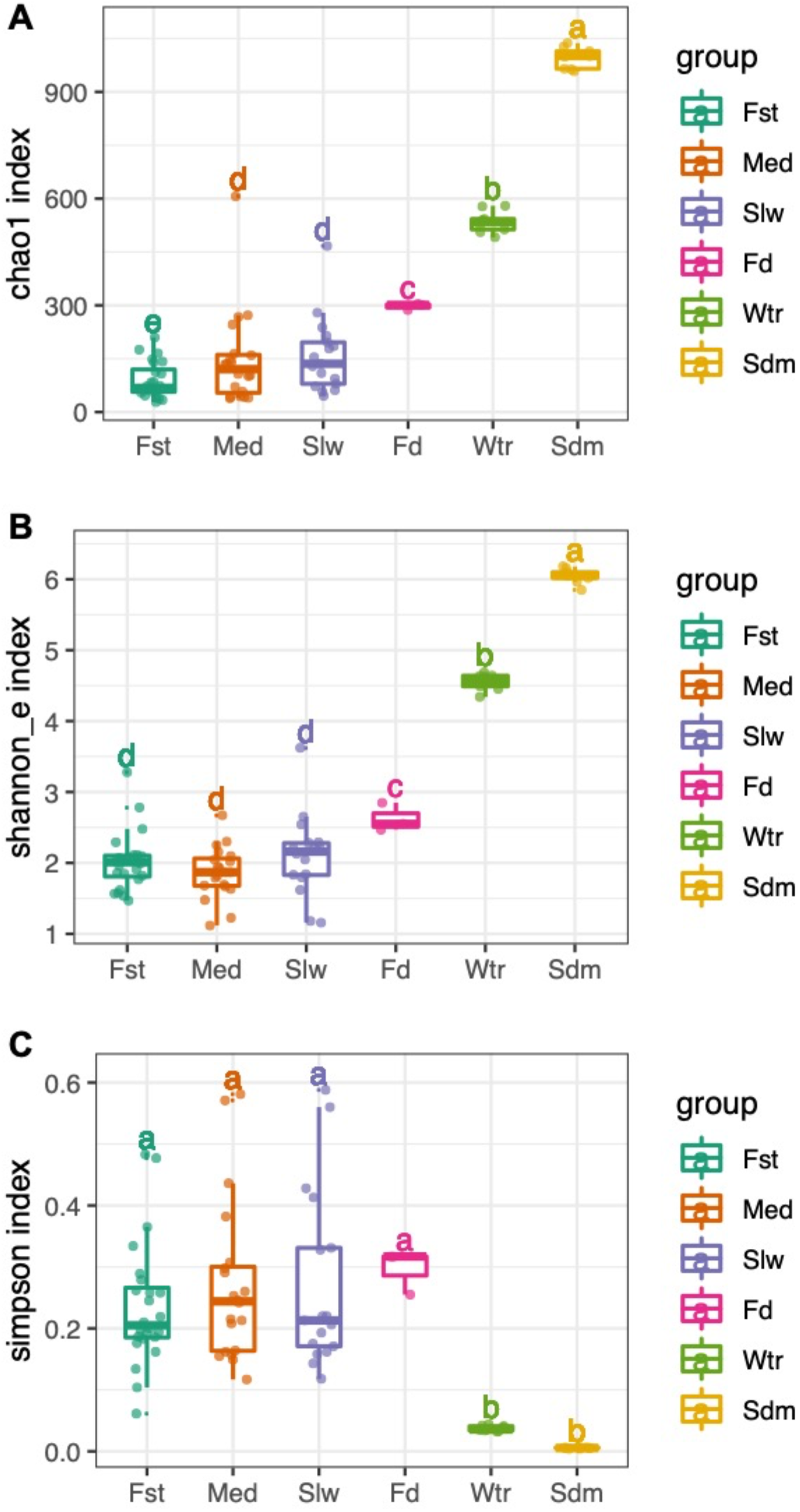
Calculated alpha-diversity based on indices of **(A)** Chao1; **(B)** Shannon_e (logarithm base *e*); **(C)** Simpson among six groups of bacterial community. For each box, the horizontal bold bar denote medians; the height of box denotes the interquartile range (25th percentile~ 75th percentile); the whiskers mark the values range within 1.5 times interquartile. Lower-cased letters denote statistical significance reported by Mann–Whitney U Test at confidence level of 0.95.

**Figure S3.**
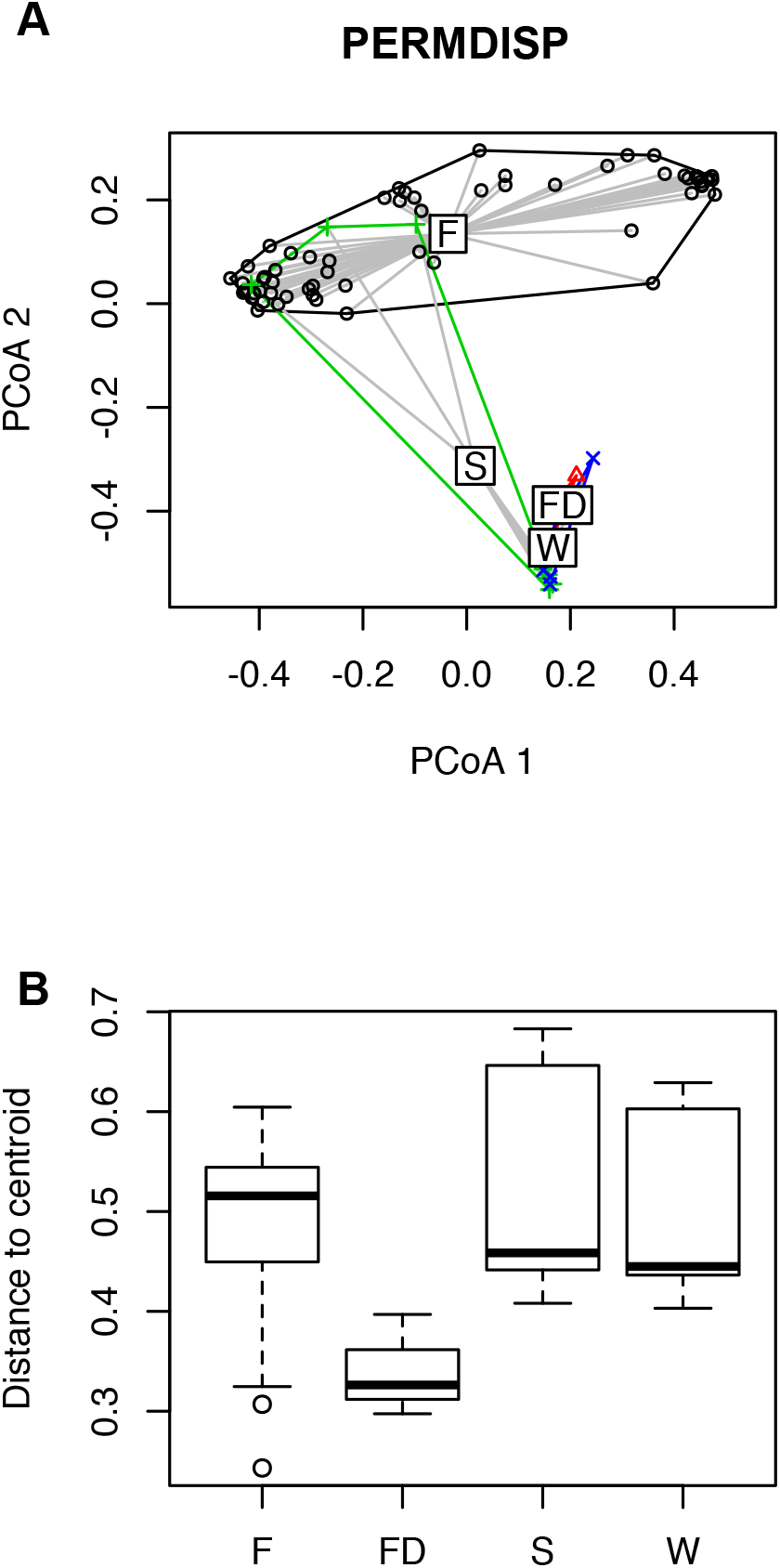
Differences in group homogeneities was examined by betadisper() implemented in R package ‘vegan’. **(A)** Principal coordinates plot showing the Bray-Curtis dissimilarity distances between each sample and its group centroids. Samples groups: F: Fugu gut; FD: feed pellet; S: sediment; W: water. **(B)** Boxplot of distances to the group centroid for the four gorups of bacterial community. For each box, the horizontal bold bar denote medians; the height of box denotes the interquartile range (25th percentile~ 75th percentile); the whiskers mark the values range within 1.5 times interquartile.

**Figure S4.**
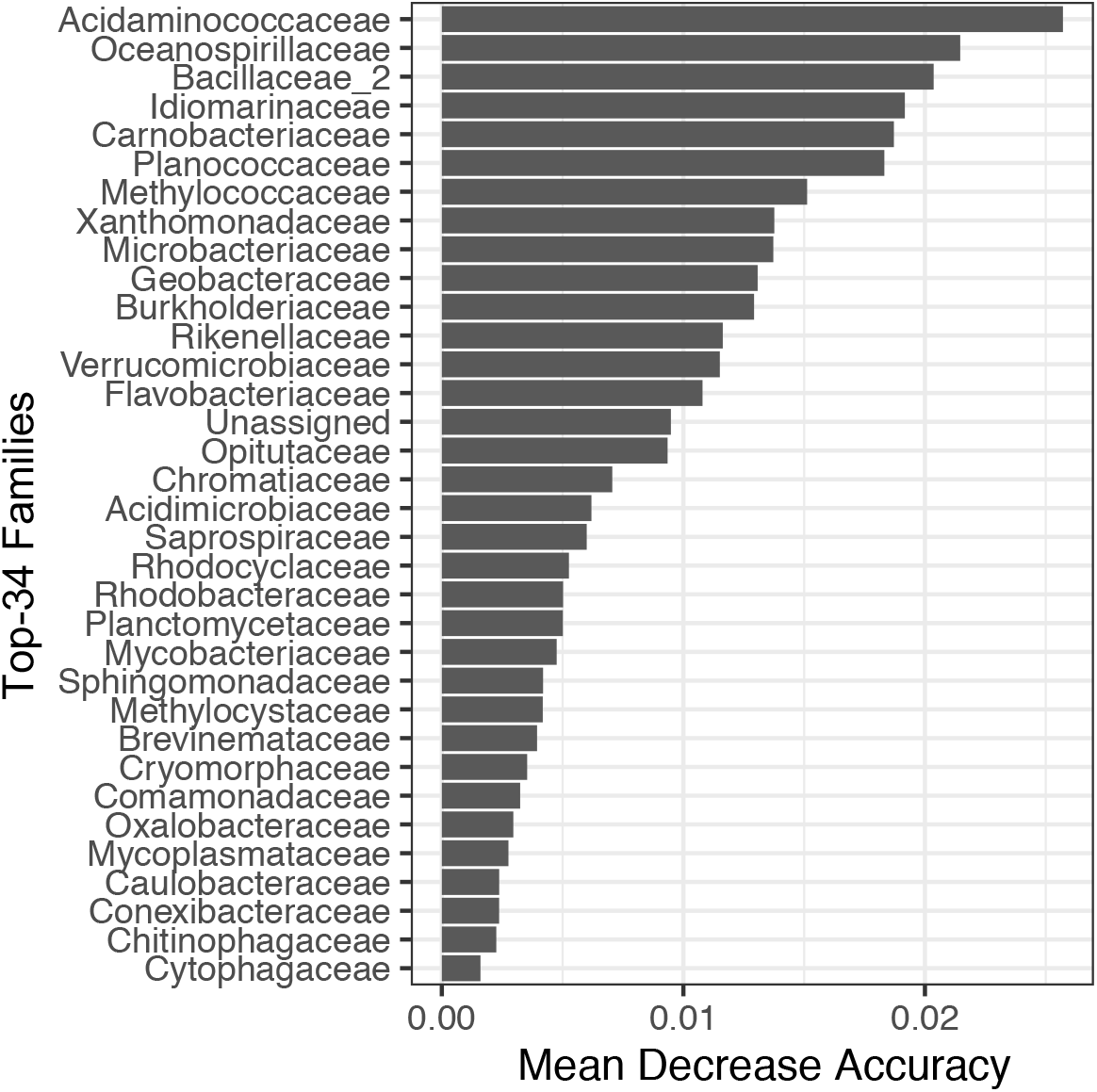
Bars of variable importance as measured by a Random Forest. The y-axis represents the top-34 bacterial families that accurately discriminated different groups of bacterial community. Bacterial families are ranked in the order of importance for group classification, from top to bottom.

**Figure S5.**
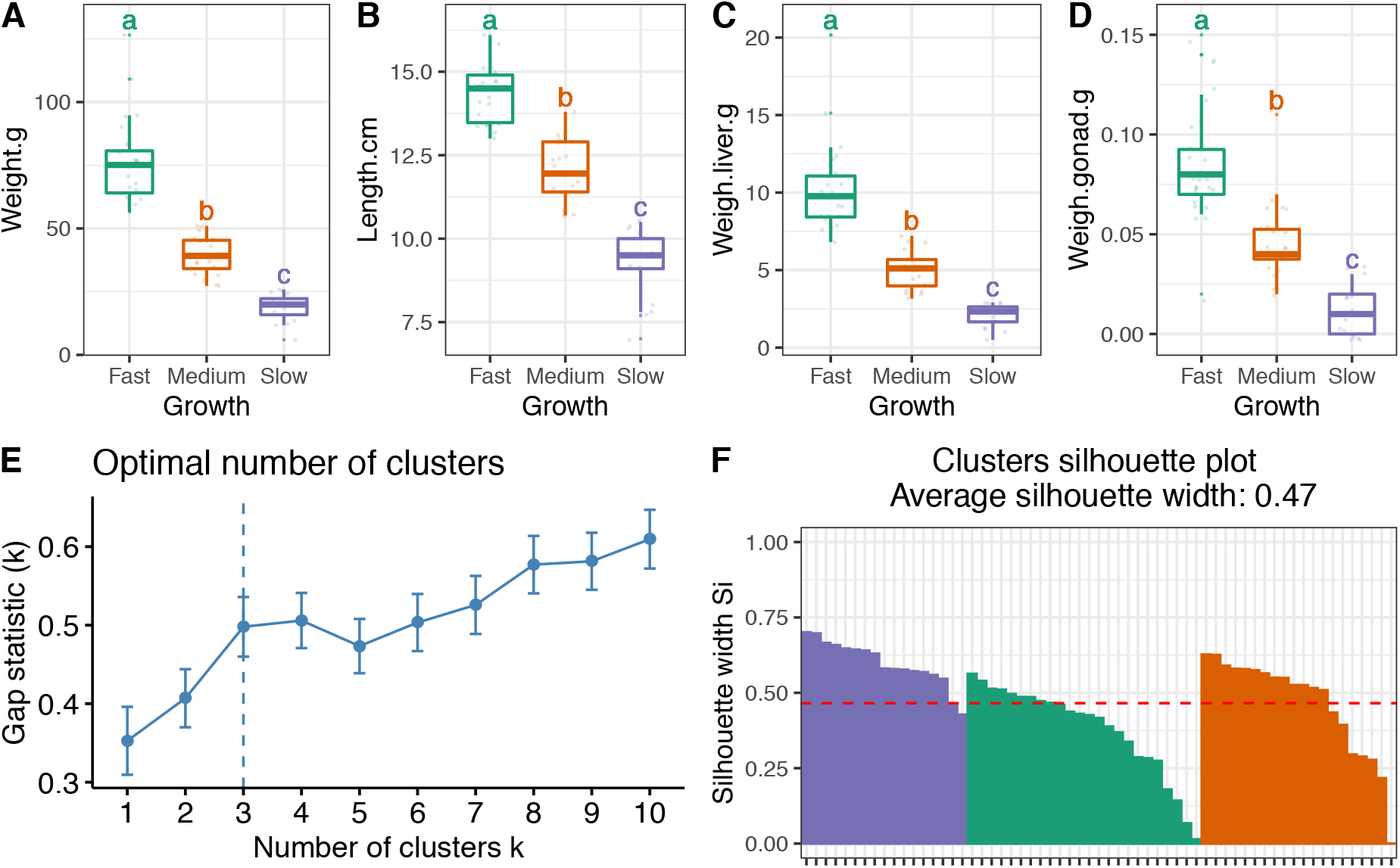
K-mean clustering based on body mass metrics. Boxplots show the differences of **(A)** body weight; **(B)** fork length; **(C)** liver weight and **(D)** gonad weight across all fish groups namely “fast”, “medium” and “**slow”.** For each box, the vertical bold bar denote medians; the width of box denotes the interquartile range (25th percentile~ 75th percentile); the whiskers mark the values range within 1.5 times interquartile. Lower-cased letters denote statistical significance reported by Mann–Whitney U test at confidence level of 0.95. **(E)** Line chart depicts the optimal numbers of clusters suggested by Gap statistic method. **(F)** Bars depecit the silhouette width for each sample, and red dash line shows the average silhouette width for k-means clustering. Bars are coloured according to the fish groups as the same in **(A~D).**

**Figure S6.**
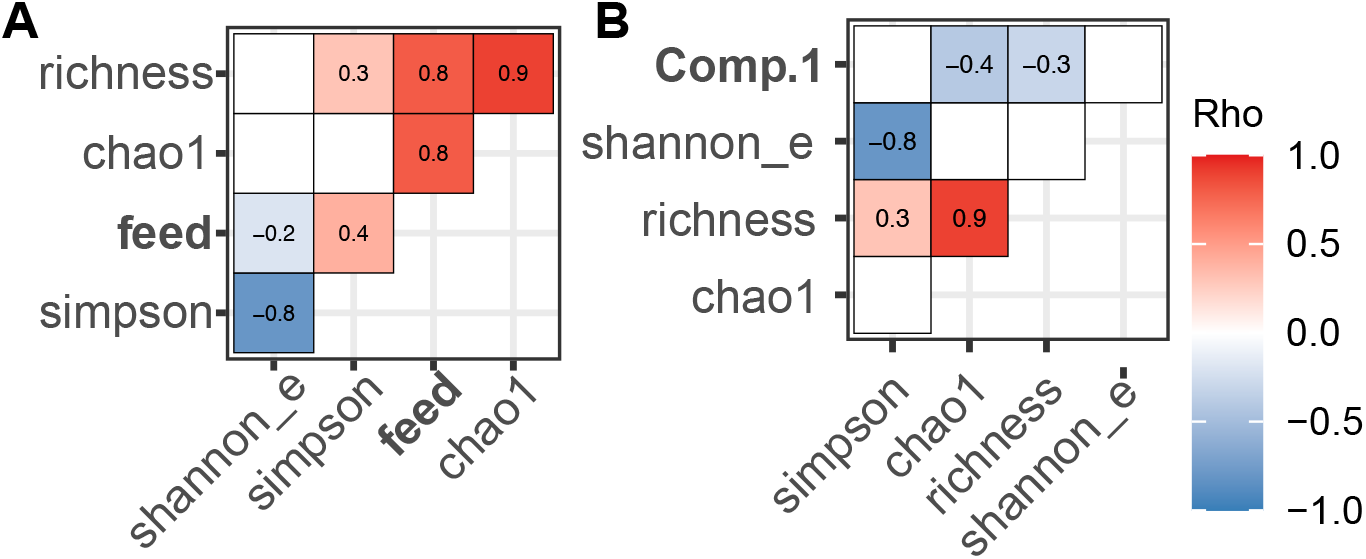
Heatmaps showing the correlations between each indicated alpha-diversity indice of fugu gut microbiota and **(A)** Feed source proportions (‘feed’) predected by SourceTracker; or **(B)** body mass indexes (‘Comp.1’) computed by PCA analysis. Color scale denotes the degree of correlatedness, only statistical significant (*P*<0.05) Spearman’s Rho were shown in each heatmap pixel.

**Figure S7.**
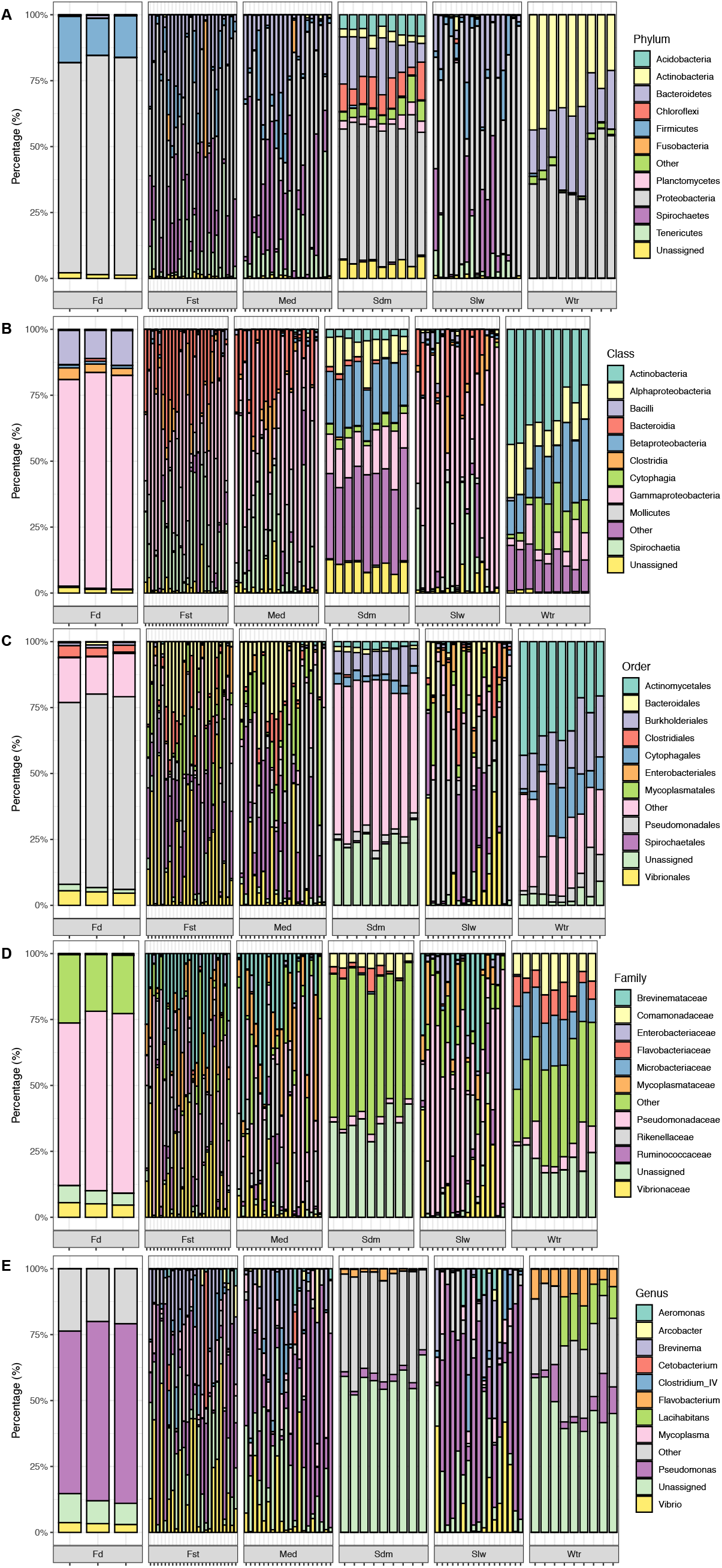
Stackbars showing the relative abundance of top-10 bacterial **(A)** phylums, **(B)** classes, **(C)** orders, **(D)** families and **(E)** genus across all biologically replicated samples. The bacterial taxonomic ranks which have lower relative abundance were grouped into “Other”.

**Figure S8.**
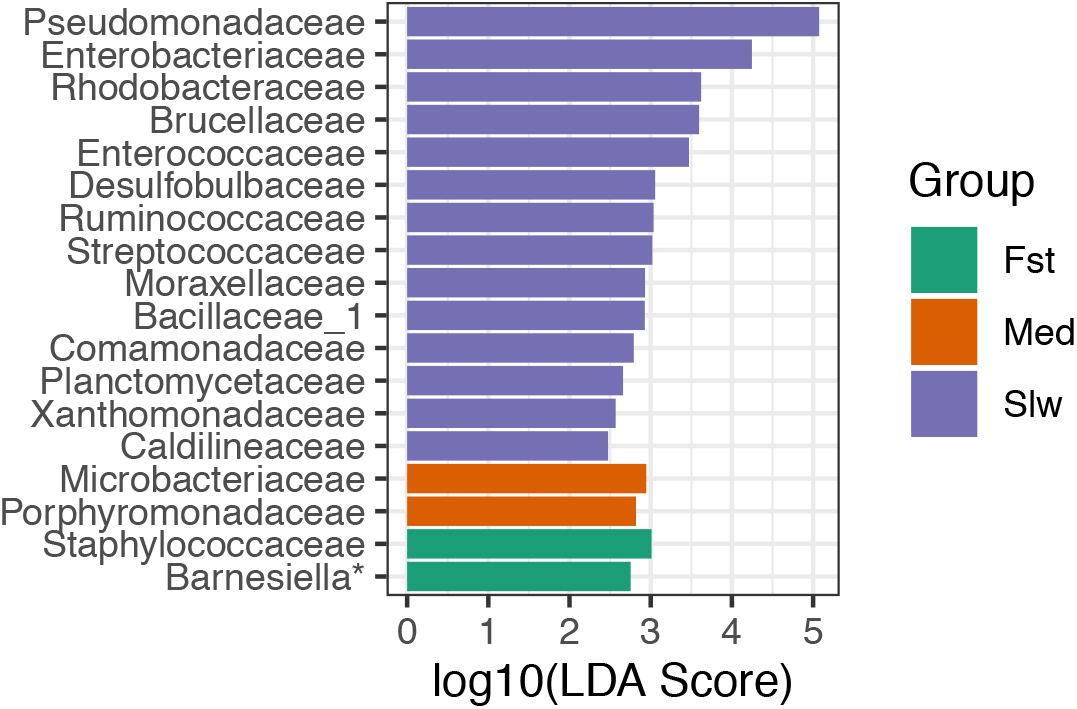
Bars show the LEfSe-estimated discriminative bacterial families that are differential among three fugu cluters with statistical significance reported by Kruskal-Wallis sum-rank test (*P*<0.05). Bar width denotes ranks of the log10-transformed LDA effect size.

**Figure S9.**
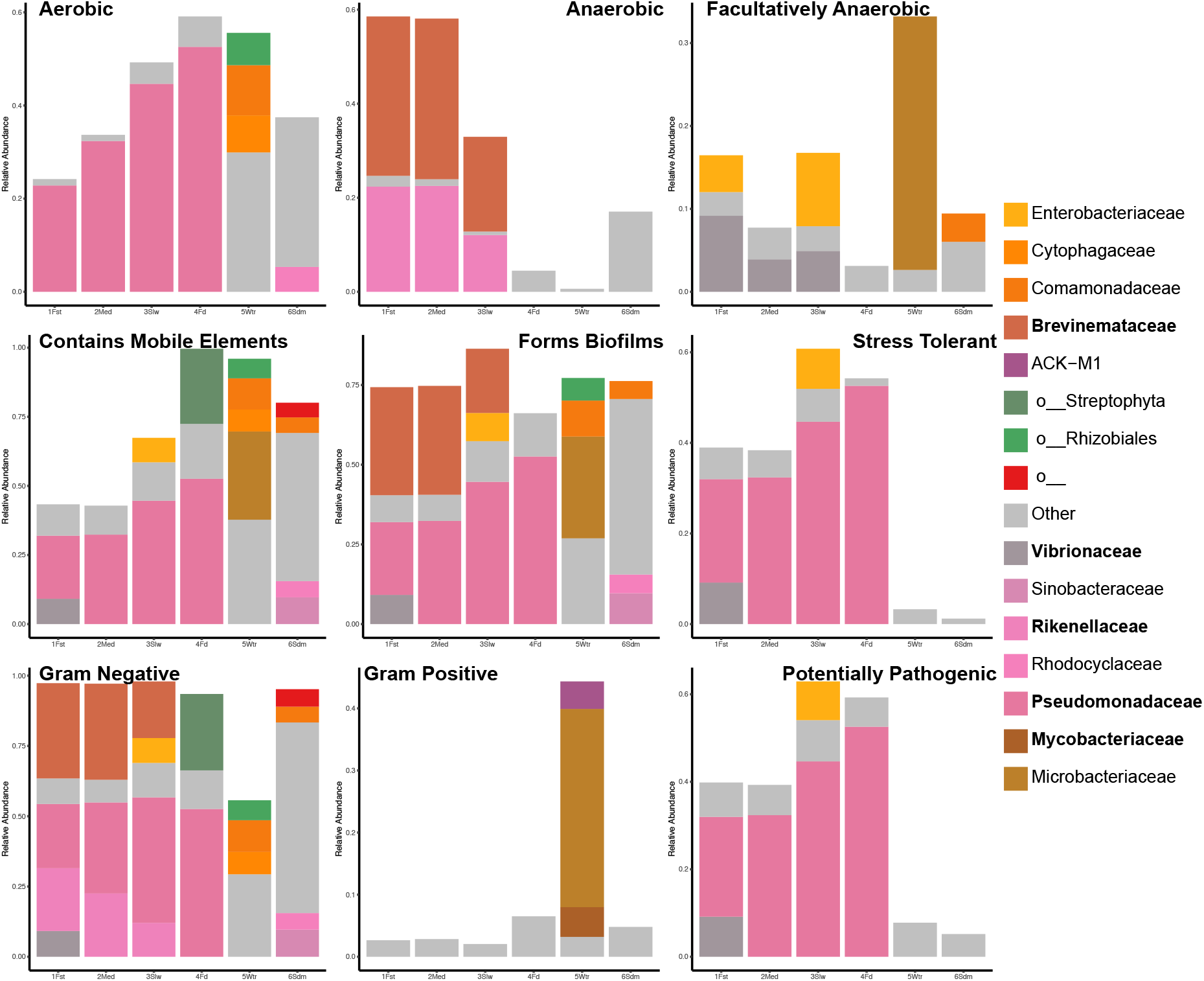
Stackbars showing the relative abundance of bacterial families in contributing to the proportions of bacterial phenotypes that are infered by BugBase. Different sample groups of bacterial communities were shown in x-axis.

**Figure S10.**
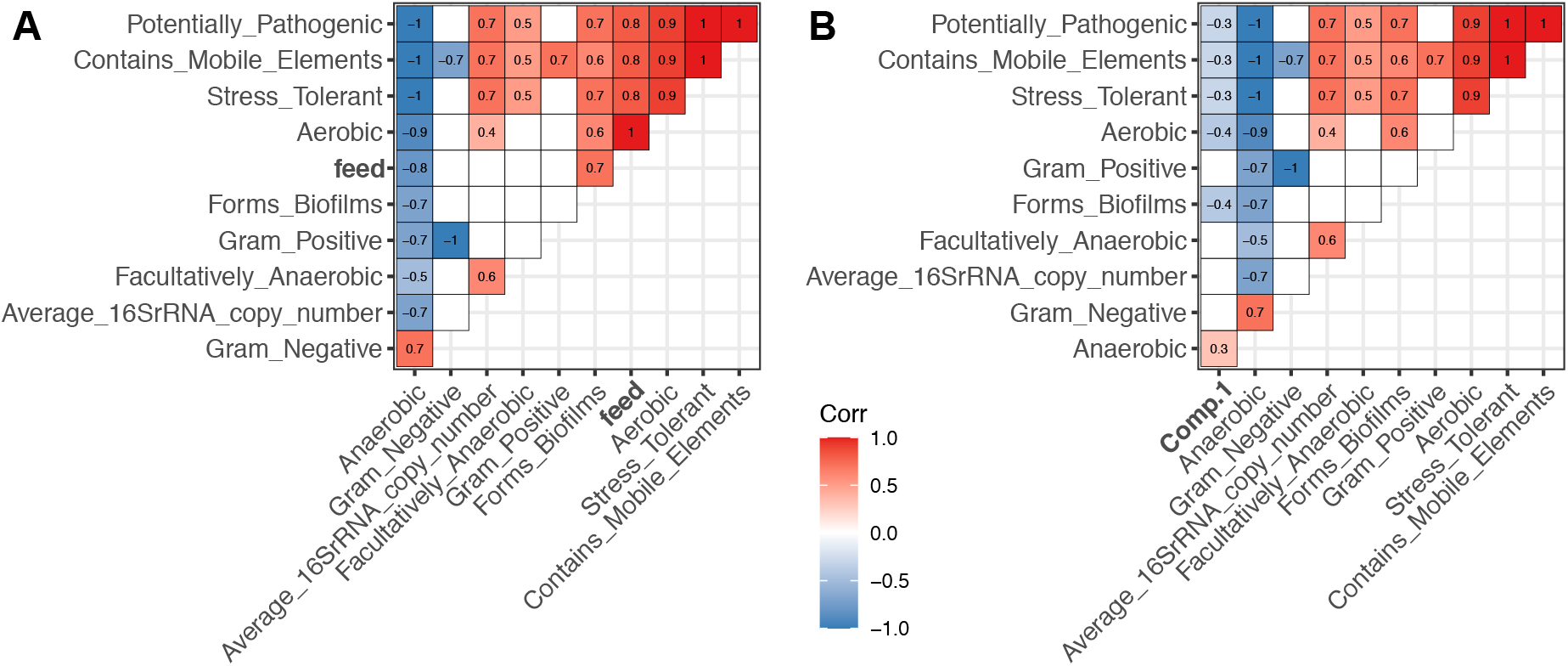
Heatmaps showing the correlations between the relative abundance of each indicated bacterial phenotype infered by BugBase and **(A)** Feed source proportions (‘feed’) predected by SourceTracker; or **(B)** body mass indexes (‘Comp.1’) computed by PCA analysis. Color scale denotes the degree of correlatedness, only statistical significant (*P*<0.05) Spearman’s Rho were shown in each heatmap pixel.

**Figure S11.**
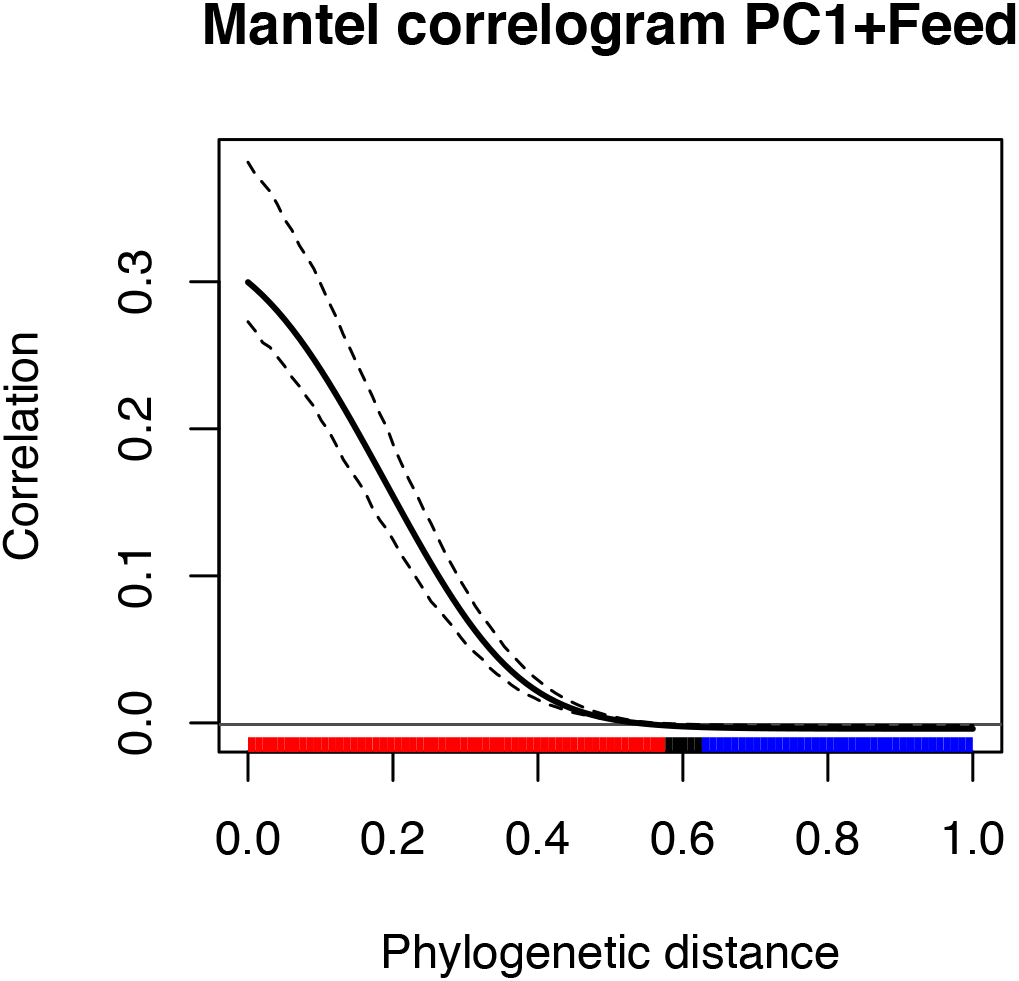
Calculation of the phylogenetic signal of traits including feed source proportions (‘feed’) predicted by SourceTracker and body mass indexes (‘PC1’) computed by PCA analysis. The solid bold black line denotes Mantel’s statistic of autocorrelation, and the dashed black lines denote the lower and upper bounds of the confidence envelop (0.95). The horizontal black line denotes the expected value of Mantel’s statistic under the null hypothesis of no phylogenetic autocorrelation. The colored bar show whether the autocorrelation is significant (based on the confidence interval): red for significant positive autocorrelation, black for nonsignificant autocorrelation, and blue for significant negative autocorrelation.

**Figure S12.**
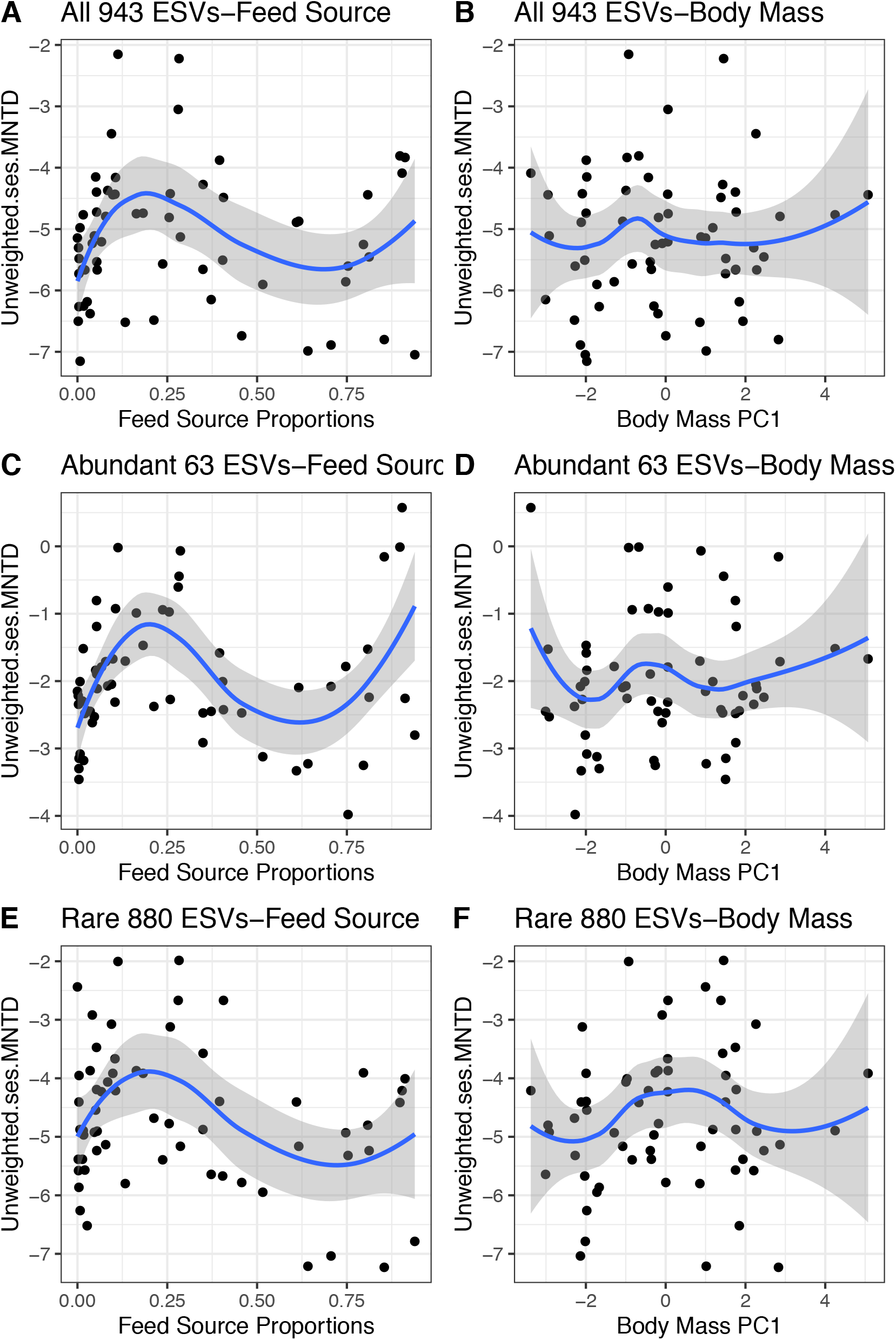
Scatterplots show the LOESS smooth curve fitting (Local Polynomial Regression) between unweighted ses.MNTD metrics of indicated sub-community with a given trait (feed source or body mass). Regressions of **(A)** feed source and **(B)** body mass with ses.MNTD for all 943 ESVs. Regressions of **(C)** feed source and **(D)** body mass with ses.MNTD for 63 abundanct ESVs. Regressions of **(E)** feed source and **(F)** body mass with ses.MNTD for 880 rare ESVs.

**Figure S13.**
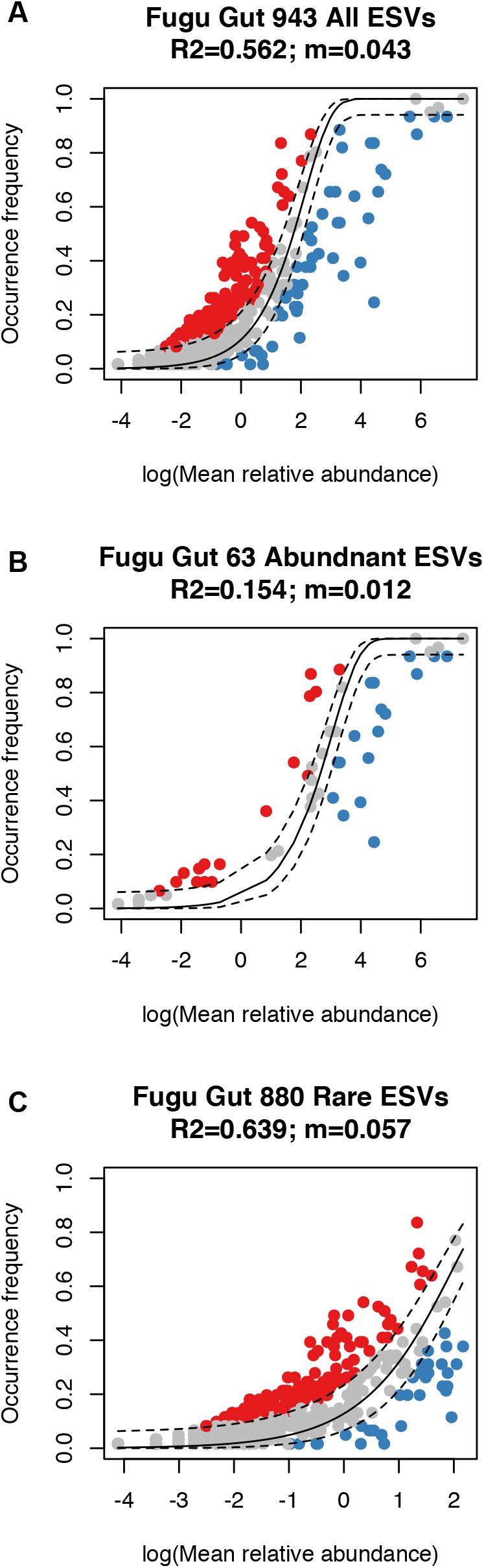
Fitness of the neutral model for **(A)** all 943 ESVs; **(B)** 63 abundanct ESVs and **(C)** 880 rare ESVs. Neutral (fit), overrepresented and underrepresented ESVs are coloured grey, red and blue respectively. The solid black line denotes model prediction, and the dashed black lines denote the lower and upper bounds of the confidence envelop (0.95). The fitness of the neutral model (R2) and migrition rate (*m*) are shown as plot title of each subpanel.

